# Targeted degradation of pathogenic TDP-43 proteins in amyotrophic lateral sclerosis using the AUTOTAC platform

**DOI:** 10.1101/2025.08.31.673396

**Authors:** Daniel Youngjae Park, Hee Yeon Kim, Eun Hye Cho, Ha Neul Choi, Da-Ha Park, Gee Eun Lee, Sung Hyun Kim, Hye Yeon Kim, Young Ho Suh, Jin-A Lee, Yong Tae Kwon

## Abstract

Amyotrophic lateral sclerosis (ALS) is a fatal neurodegenerative disease characterized by the progressive loss of motor neurons and the cytoplasmic aggregation of misfolded proteins in the spinal cord, including TAR DNA-binding protein-43 (TDP-43). More than 97% of ALS cases exhibit pathological TDP-43 inclusions, yet therapeutic strategies that can selectively eliminate these aggregates remain yet to be developed. Here, we employed the AUTOTAC (Autophagy-Targeting Chimera) to degrade TDP-43 aggregates via macroautophagy mediated by the N-recognin p62/SQSTM1 of the N-degron pathway. The AUTOTAC degraders ATC141 and ATC142 were designed to bind and link the oligomeric species of misfolded TDP-43 to p62, which induces the targeting of TDP-43 cargoes to phagophores for lysosomal co-degradation, while sparing monomeric TDP-43. ATC142 induced the degradation of pathological TDP-43 A315T species and its cleaved variant, TDP-25, with DC_50_ values of 1.25-9.6 nM. In ALS model mice expressing TDP-43 A315T in the spinal cord, oral administration of 10 mg/kg ATC141 with 24 doses reduced TDP-43 aggregates as well as GFAP^+^ astrocytes and Iba1^+^ microglia. ATC141 also exerted disease-modifying efficacy to reverse the disease progression in neuromuscular coordination and cognitive function. This oral drug is under Phase 1 clinical trials in South Korea with 76 healthy volunteers aiming to treat ALS, Alzheimer’s diseases (AD), and progressive supranuclear palsy (PSP). We suggest that AUTOTAC provides a novel strategy to treat a broad range of neurodegenerative diseases.

**Teaser:** AUTOTAC degraders induce lysosomal degradation of pathogenic TDP-43 aggregates in a mouse model of amyotrophic lateral sclerosis.

## Introduction

ALS is a fatal neurodegenerative disease characterized by the progressive degeneration of upper and lower motor neurons in the brain and spinal cord (*1*). The high mortality associated with ALS is driven by progressive muscle paralysis, ultimately resulting in death due to respiratory failure within 3-5 years of diagnosis (*2*). Approximately 90-95% of ALS cases are sporadic with no apparent history of the disorder in the family (*3*). Sporadic ALS (sALS) arises from heterogenous factors, primarily oxidative stress and mitochondrial dysfunction, and is characterized by progressive muscle weakness and atrophy that spread throughout the body (*4*). Processes such as glutamate excitotoxicity or glial activation alone are not able to induce motor neuron damage or drive ALS progression (*5, 6*), and are likely a secondary response. During the disease progression, ALS conditions stimulate the aggregation of specific cellular proteins such as TAR DNA-binding protein (TDP-43), which further exacerbates neuroinflammation, synaptic loss, and motor neuron degeneration, ultimately leading to cell death (*7*). In contrast to sALS, approximately 5-10% of ALS cases have a family history of ALS or frontotemporal dementia (FTD) that progressively affects personality, behavior, and language (*8*). Familial ALS (fALS) is associated with mutations in more than 25 genes, including *C9ORF72*, *SOD1*, *TARDBP*, *FUS*, *VCP*, *OPTN*, and *TBK1* (*9*). Amongst these, repeat expansions in *C9ORF72* represent approximately 40% of fALS, contributing to the formation of toxic RNA foci, dipeptide repeat (DPR) proteins, and insoluble aggregates (*10*). Mutations in superoxide dismutase 1 (SOD1) and *TARDBP*-encoded TDP-43 accounts for about 10% and 4% of fALS, respectively, and commonly lead to their misfolding and aggregation that trigger cellular toxicity (*11*).

TDP-43 is a nuclear protein that regulates the transcription, splicing, and stability of RNA (*12*). During central nervous system (CNS) development in early embryos, TDP-43 is highly expressed in the neuroepithelium, the source of CNS progenitor cells (*13*). Under ALS conditions characterized by dysregulated reactive oxygen species (ROS) and impaired nucleocytoplasmic transport, 50-80% of nuclear TDP-43 is mislocalized to the cytoplasm (*14*). Following relocalization, cytosolic TDP-43 undergoes abnormal post-translational modifications, which induce its destabilization and misfolding (*15*). Misfolded TDP-43 can be cleaved into toxic C-terminal fragments, as represented by TDP-25, which synergistically accelerates the formation of insoluble, toxic aggregates (*16*). Furthermore, TDP-43 acts as a nucleating seed that synergistically drives the misfolding and co-aggregation of other ALS-associated proteins (*17*). The misfolding of wild-type SOD1 in sALS, for instance, is induced by mislocalized TDP-43, resulting in the accumulation of SOD1-positive inclusions in non-SOD1 ALS cases (*18*). Through cross seeding and co-aggregation of ALS proteins, the misfolded aggregates of wild-type TDP-43 are found in approximately 97% of sALS cases (*19*). This mechanism of TDP-43 proteinopathies also underly in other neurodegenerative disorders, including AD where TDP-43 co-aggregates with misfolded tau in up to 57% of cases (*20*). In Parkinson’s disease (PD), the co-aggregation of TDP-43 and α-synuclein through their cross-seeding exacerbates the neurotoxicity of dopaminergic neurons in approximately 7% of PD cases (*17*). In frontotemporal dementia (FTD), TDP-43 pathology is a hallmark present in ∼50% of cases, where it disrupts RNA processing, impairs synaptic integrity and plasticity, and contributes to chronic neuroinflammation (*21*). Thus, TDP-43 proteinopathy is implicated across a broad spectrum of neurodegenerative diseases, either as a central or contributing feature to disease progression. Given the aggregation-prone nature of both wild-type and mutant TDP-43, along with its prion-like propagation and cross-seeding abilities, its aggregates are emerging as a key therapeutic target across a range of neurodegenerative disorders.

For the past decades, extensive efforts have been made to develop therapeutics capable of treating ALS. The therapeutic targets available in the market include mitochondria, oxidative stress, ferroptosis, hypoxia, and inflammation (*22*). Riluzole, an inhibitor of glutamatergic neurotransmission, was approved by the US FDA in 1995 to treat ALS (*23*). However, its efficacy is modest, prolonging tracheostomy-free survival by only 2-3 months (*23*). Edaravone was approved to scavenge ROS in ALS patients but did not show significant clinical benefit in a long-term study (*24*). Tofersen, an antisense oligonucleotide (ASO) targets SOD1 mRNA in ALS patients carrying SOD1 mutations (*25*). While tofersen has shown promise in lowering SOD1 protein levels and improving biomarker profiles, its clinical impact on disease progression remains unclear. As an alternative approach, recent studies employed targeted protein degradation (TPD) to eliminate ALS-associated proteins (*26*). Amongst TPD technologies, PROTAC (proteolysis-targeting chimera) uses heterobifunctional small molecules that recruit an E3 ligase to the target substrate (*27*). A PROTAC designed to selectively degrade C-terminal fragments of TDP-43 (C-TDP-43) aggregates improved motility in a *Caenorhabditis elegans* ALS model (*28*). However, given the inabilities of the chaperones and the proteasome to unfold TDP-43 oligomers for degradation and the inner diameter of the proteasome acting as a physical barrier (*28, 29*) there are significant questions on how these TPD strategies would tackle oligomeric, fibrillar, or larger aggregates that are sized beyond the capacities of the aforementioned degradation systems. Hence, autophagy-dependent TPD technologies are a valuable tool for the removal of large protein aggregates, since the aggregates have been untouchable by the other TPD technologies.

The disturbance in proteostasis is a key pathological process in neurodegenerative diseases (*30*). Protein homeostasis depends on a protein quality control (PQC) system to ensure timely degradation of misfolded proteins, preventing their accumulation into energetically favorable insoluble aggregates (*31, 32*). These misfolded proteins are primarily degraded by the ubiquitin-proteasome system (UPS), which serves as the first line of defense in proteostasis (*31*). While efficient in eliminating smaller, short-lived soluble proteins, the capacity of UPS to degrade larger or aggregated proteins is limited (*33*). Larger oligomeric forms resistant to the UPS requires the activity of autophagy for their clearance (*34*). In ALS, the accumulation of aggregated TDP-43 saturates the proteasome’s capacity, leading to a compensatory reliance on autophagy for degradation (*35*). Moreover, TDP-43 aggregates are known to physically block autophagic structures or sequester essential autophagy proteins, further impairing autophagic pathways (*36*). Neurons are particularly vulnerable to proteostasis disruptions due to their unique biological constraints including their post-mitotic nature and high metabolic demand (*37*). These cumulative proteostasis impairments create a cellular environment where misfolded and aggregated proteins persist, exacerbating neuronal dysfunction and degeneration. To counteract, restoring basal autophagy or enhancing lysosomal capacity may offer potential therapeutic strategies.

The N-degron pathway is a degradative system in which specific N-terminal residues, termed N-degrons, are recognized as signals to induce substrate degradation (*38, 39*). N-degrons, such as N-terminal arginine (Nt-Arg), are selectively recognized by N-recognins including E3 ligases of the UBR subfamily, which leads to ubiquitination and subsequent proteasomal degradation (*40–42*). We have previously identified the autophagic N-degron pathway in which autophagy receptor p62/SQSTM1 acts as an N-recognin that binds Nt-Arg via its ZZ-domain (*43*). The N-degron Nt-Arg can be generated by ATE1 R-transferases that conjugate the amino acid L-Arg from Arg-tRNA^Arg^ to protein N-termini (*44*). Upon binding with Nt-Arg, p62 undergoes a conformational change, exposing its PB1 domain, which facilitates self-polymerization in complex with its cargoes, such as protein aggregates (*43, 45, 46*). Simultaneously, the exposed LIR domain of Nt-Arg bound p62 interacts with LC3 on phagophores, leading to lysosomal degradation of p62-cargo complexes (*46, 47*). This mechanism-of-action of the Arg/N-degron pathway mediates degradation of misfolded protein aggregates and subcellular organelles such as the endoplasmic reticulum (ER) and peroxisomes via p62-dependent autophagy (*48–52*).

To harness the N-degron pathway for therapeutic applications, we developed chemical mimetics of N-degrons, termed autophagy-targeting ligands (ATLs) that target p62 ZZ domain, mimicking the biological function of N-degrons to activate autophagy (*43*). Based on the ATL technology, we also developed AUTOTAC that enabled targeted degradation of proteins via p62-mediated autophagy (*43, 46*). AUTOTAC employs a chimeric configuration composed on a target-binding ligand (TBL) linked to an ATL that binds and activates p62 via N-degron interactions (*46*). Compared to PROTACs, AUTOTAC leverages the autophagy-lysosomal pathway, enabling the degradation of larger and more stable aggregates, including oligomeric species of aggregation-prone proteins in neurodegenerative diseases.

In this study, we designed and synthesized AUTOTACs to selectively target and degrade pathological TDP-43 oligomers, thereby reducing protein aggregation. Under ALS conditions, TDP-43 misfolds primarily within its aggregation-prone C-terminal domain, also referred to as the prion-like domain, where normally buried hydrophobic patches around residues 320-340 and 370-390 become exposed (*53, 54*). This conformational change drives β-sheet-rich oligomerization and the formation of amorphous aggregates (*54*). By binding to β-sheet-rich oligomeric interfaces, ATC141 and ATC142 induce p62-dependent targeted autophagic-lysosomal degradation, thereby mitigated hyperactivated NF-κB p65 signaling. Orally administrable ATC141 effectively cleared TDP-43 aggregates in ALS patient-derived induced pluripotent stem cell (iPSC) motor neurons and ALS-afflicted mice, ultimately reducing glial activation and improving motor function. In summary, this study highlights targeted degradation of pathogenic TDP-43 species as a promising therapeutic strategy for ALS, emphasizing AUTOTACs’ potential as a first-in-class therapeutic agent.

## Results

### Development of the AUTOTAC ATC142 that induces the degradation of pathogenic TDP-43 species via p62-mediated macroautophagy

Approximately 50-60% of TDP-43 is composed of intrinsically disordered regions (IDRs), making it difficult to design small molecule ligands with high selectivity (*55*). We therefore aimed to design an AUTOTAC whose TBL recognizes the β-sheet structure shared by pathogenic TDP-43 species in their oligomeric and fibrillar states. Through the screening of a set of AUTOTACs carrying different TBLs, including Anle138b, 4-phenylbutyrate (4-PBA), curcumin, and methylene blue (fig. S1A), we further characterized ATC142 harboring a derivative of Anle138b as a TBL (fig. S1B, S2, and fig. S3, A to D). In silico binding analysis between Anle138b and the amyloidogenic human TDP-43 structure (PDB ID: 7PY2) revealed a stable interaction within residues 311-360, the fibrillary core region. Molecular docking showed a Glide SP score of −4.293 and a favorable MM-GBSA binding free energy of −65.4 kcal/mol (table S1). The 2D interaction diagram (Fig. 1A) highlighted extensive contacts, including hydrogen bonds with G335 and N343 (Fig. 1B), supporting the use of the Anle138b scaffold for selective recognition of β-sheet-rich TDP-43 conformers in their oligomeric and fibrillary states. To assess the degradation efficacy of ATC142, we used HEK293T and HeLa cells transiently expressing TDP-43 A315T mutant known to undergo spontaneous misfolding and aggregation (*56*). ATC142 exhibited a DC_50_ (half-maximal degradation concentration) for TDP-43 A315T tagged with GFP at the range of 1.25-25 nM in a dose-dependent manner with D_max_ (maximal degradation efficacy) of approximately 90% (Fig. 1, C and D, and fig. S3, A and B). As an alternative model substrate, we also used TDP-25, a highly aggregation-prone and neurotoxic fragment. The DC_50_ of TDP-25 ranged from 1.0 to 9.6 nM with the D_max_ of over 90% in HEK293T and HeLa cells (Fig. 1, E and F and fig. S3, C and D). Similar degradation efficacy was observed with TDP-43 A315T tagged with flag and TDP-25 tagged with myc (Fig. 1G and fig. S3E). The T_1/2_ (time for 50% degradation) was estimated to be 35 and 20 min, respectively, for TDP-43 A315T and TDP-25 (Fig. 1, H to K). In washout assay with the proteasome inhibitor MG132, the degradation efficacy of ATC142 persisted up to 8 hours, followed by a hook effect (Fig. 1L). In contrast to ATC142, no significant degradation was observed with the ATL and TBL alone (fig. S3, F to K). These results indicate that ATC142 induces the degradation for pathological TDP-43 species, possibly through a catalytic cycle.

**Fig. 1.**
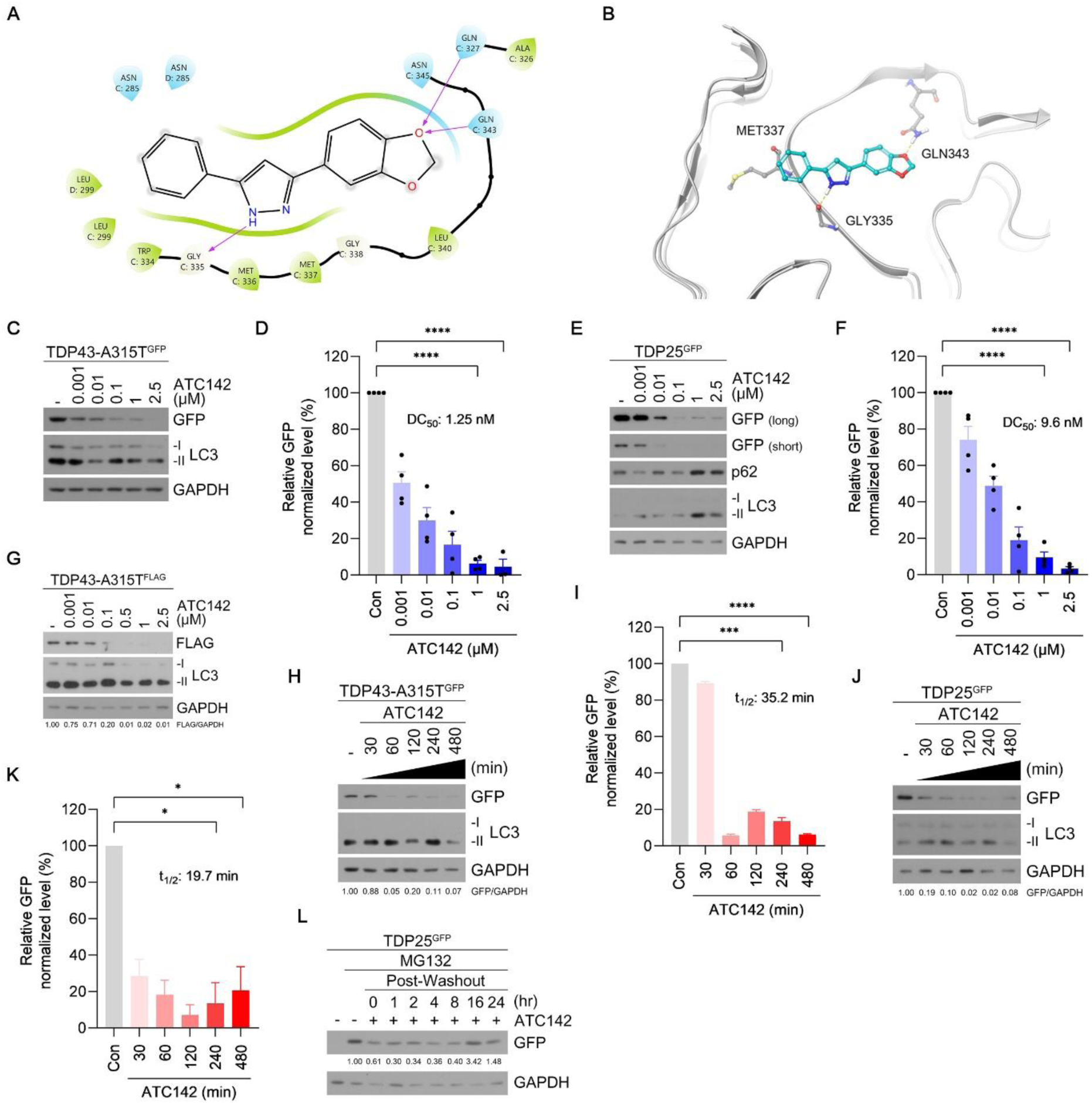
Targeted degradation of pathogenic TDP-43 species by ATC142. **(A)** Schematic representation of the 2D ligand interaction diagram illustrating multiple contacts between Anle138b and the amyloidogenic human TDP-43 structure (PDB ID: 7PY2). Hydrogen bond interactions are shown as purples lines, and π-cation interactions as red lines. **(B)** The 3D molecular docking pose of Anle138b (cyan) bound to amyloidogenic TDP-43 (grey). Hydrogen bonds are represented as green dotted lines. **(C)** Immunoblotting analysis of HeLa cells transfected with TDP-43 A315T-GFP and treated with ATC142 at the indicated concentrations (24 h). **(D)** Quantification of GFP levels shown in (C). **(E)** Identical to (C), but in HeLa cells transfected with TDP25-GFP. **(F)** Quantification of GFP levels shown in (E). **(G)** Immunoblotting analysis of HeLa cells transfected with TDP-43 A315T-FLAG and treated with ATC142 at the indicated concentrations (24 h). **(H)** Immunoblotting analysis of HeLa cells transfected with TDP-43 A315T-GFP and treated with ATC142 (1 µM) for the indicated durations. **(I)** Quantification of GFP levels shown in (H). **(J)** Identical to (H), but in HeLa cells transfected with TDP25-GFP. **(K)** Quantification of GFP levels shown in (J). **(L)** Immunoblotting analysis of HeLa cells transfected with TDP25-GFP and treated with MG132 (1 µM, 24 h) and ATC142 (1 µM, 24 h), followed by the culture medium subsequently washed out for the indicated durations. Data are presented as mean ± S.E.M. Statistical significance is indicated as follows: *P ≤ 0.05; ***P ≤ 0.001; ****P ≤ 0.0001. (Student’s *t*-test).

The pathogenesis of ALS involves the cytosolic mislocalization of nuclear TDP-43 and its misfolding and cleavage into TDP-25 (*19*). We therefore determined whether ATC142 enables p62 to hijack cytosolic TDP-43 A315T species for lysosomal degradation. Knockdown of p62 using siRNAs abolished ATC142-induced degradation of TDP-43 A315T and TDP-25, resulting in a marked accumulation of both species (Fig. 2, A and B). In co-immunoprecipitation (co-IP) analysis, ATC142 enhanced the interaction of TDP-25 with p62 (Fig. 2C) but not with its ZZΔ mutant (Fig. 2D). When autophagic flux was blocked using hydroxychloroquine (HCQ) that inhibits lysosomal acidification, ATC142 induced the accumulation of TDP-43 as well as the lipidation of LC3, which were degraded by lysosomal hydrolases (Fig. 2, E and H). Autophagy flux assays using HCQ followed by immunocytochemistry (ICC) revealed that ATC142 induced the formation of cytosolic puncta positive for p62 and TDP-43 A315T (Fig. 2, F and G). ATC142-dependent formation of p62-associated complexes destined to lysosomal degradation was similarly observed with TDP-25 (Fig. 2, I and J). The majority of p62^+^TDP-43 A315T^+^ and p62^+^TDP25^+^ cytosolic puncta induced by ATC142 were also positive for LC3, indicative of autophagic sequestration (Fig. 2, K and L). Consistently, knockdown of ATG5 or ATG7, essential for autophagy initiation and autophagosome biogenesis, respectively, suppressed ATC142-mediated degradation of TDP-25 (Fig. 2, M and N). Ubiquitin depletion using *Ubb* siRNA did not significantly impacted the degradation of both TDP-43 A315T and TDP-25 induced by ATC142 (Fig. 2, O and P). These results collectively demonstrate that ATC142 targets pathological TDP-43 aggregates for degradation via p62-dependent, ubiquitin-independent autophagy.

**Fig. 2.**
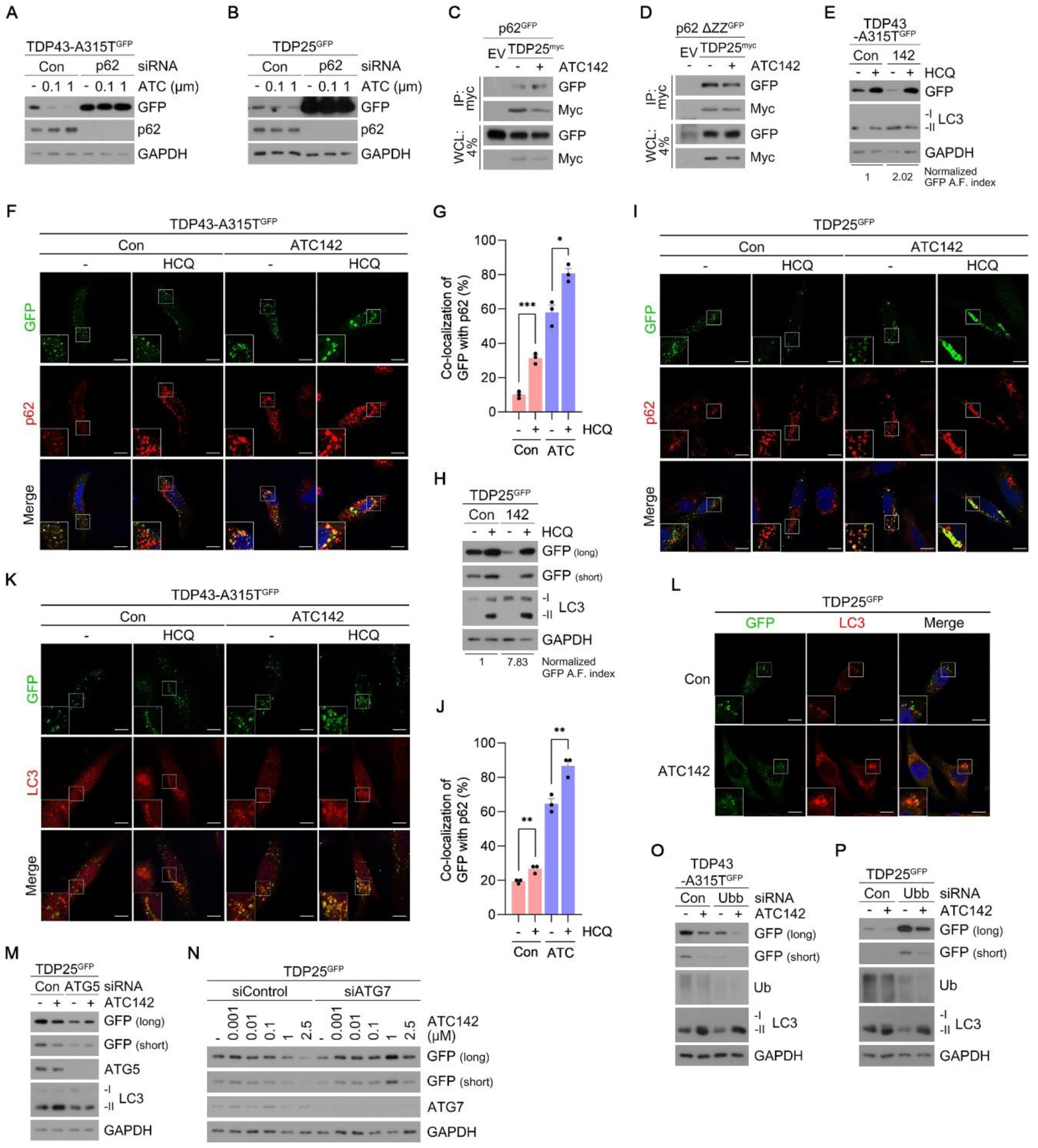
ATC142 targets TDP-43 aggregates to p62-mediated macroautophagy. **(A)** Immunoblotting analysis of HeLa cells transfected with TDP-43 A315T-GFP and treated with ATC142 at the indicated concentrations (24 h) following RNA interference of p62 (40 nM, 24 h). **(B)** Identical to (A), but in HeLa cells transfected with TDP25-GFP. **(C)** Co-immunoprecipitation assay in HEK293T cells co-expressing p62-GFP and TDP25-Myc, treated with ATC142 (1 µM, 24 h). **(D)** Identical to (C), but in HEK293T cells co-expressing p62△ZZ-GFP and TDP25-Myc. **(E)** Autophagic flux assay in HeLa cells transfected with TDP-43 A315T-GFP treated with ATC142 (1 µM, 24 h), in the presence or absence of HCQ (10 µM, 24 h). **(F)** Immunocytochemical detection of p62 in HeLa cells transfected with TDP-43 A315T-GFP and treated with ATC142 (1 µM, 24 h), in the presence or absence of HCQ (10 µM, 24 h). Scale bar, 10 µm. **(G)** Quantification of GFP and p62 co-localization shown in (F). **(H)** Identical to E, but in HeLa cells transfected with TDP25-GFP. **(I)** Identical to F, but in HeLa cells transfected with TDP25-GFP. Scale bar, 10 µm. **(J)** Quantification of GFP and p62 co-localization shown in (I). **(K)** Immunocytochemical detection of LC3 in HeLa cells transfected with TDP-43 A315T-GFP and treated with ATC142 (1 µM, 24 h), in the presence or absence of HCQ (10 µM, 24 h). Scale bar, 10 µm. **(L)** Immunocytochemical detection of LC3 in HeLa cells transfected with TDP25-GFP and treated with ATC142 (1 µM, 24 h). Scale bar, 10 µm. **(M)** Immunoblotting analysis of HeLa cells transfected with TDP25-GFP and treated with ATC142 (1 µM, 24 h) following RNA interference of ATG5 (40 nM, 48 h). **(N)** Identical to (M), but following RNA interference of ATG7 (40 nM, 48 h). **(O)** Immunoblotting analysis of HeLa cells transfected with TDP-43 A315T-GFP and treated with ATC142 (1 µM, 24 h) following RNA interference of *Ubb* (40 nM, 48 h). **(P)** Identical to (O), but in HeLa cells transfected with TDP25-GFP. Data are presented as mean ± S.E.M. Statistical significance is indicated as follows: *P ≤ 0.05; **P ≤ 0.01; ***P ≤ 0.001. (Student’s *t*-test).

### ATC142 selectively eliminates aggregated species of TDP-43 while preserving its functional monomers

Cytosolic relocalization of TDP-43 in degenerating neurons leads to it aggregation (*19, 57*). To assess the selectivity of ATC142 to oligomeric and aggregated species of misfolded TDP proteins, TDP-25 (fig. S4A) or TDP-43 A315T (Fig. 3A, lane 9) was transiently expressed in HEK293 cells to induce cytosolic accumulation of their high molecular-weight species. When cells were fractionated, both TDP-25 and TDP-43 A315T prominently relocated to the cytosol to form SDS-resistant aggregates, which were efficiently degraded following ATC142 treatment (Fig. 3A, lane 10 and Fig. 3B). By sharp contrast, ATC142 induced no detectible degradation for their nuclear species (Fig. 3B lane 4). Next, to further accelerate the cytosolic relocation and aggregation of TDP-43 and its fragments, HEK293 cells expressing TDP-25 or TDP-43 A315T were treated with sodium arsenite, a metalloid compound that induces oxidative stress and cellular damage. Non-reducing SDS-PAGE showed that ATC142 reduced the levels of their oligomeric and aggregated species but not monomers (Fig. 3, C and D). These results indicate that ATC142 induces the degradation of aggregated species of TDP-43 and TDP-25 but not their functional monomers.

**Fig. 3.**
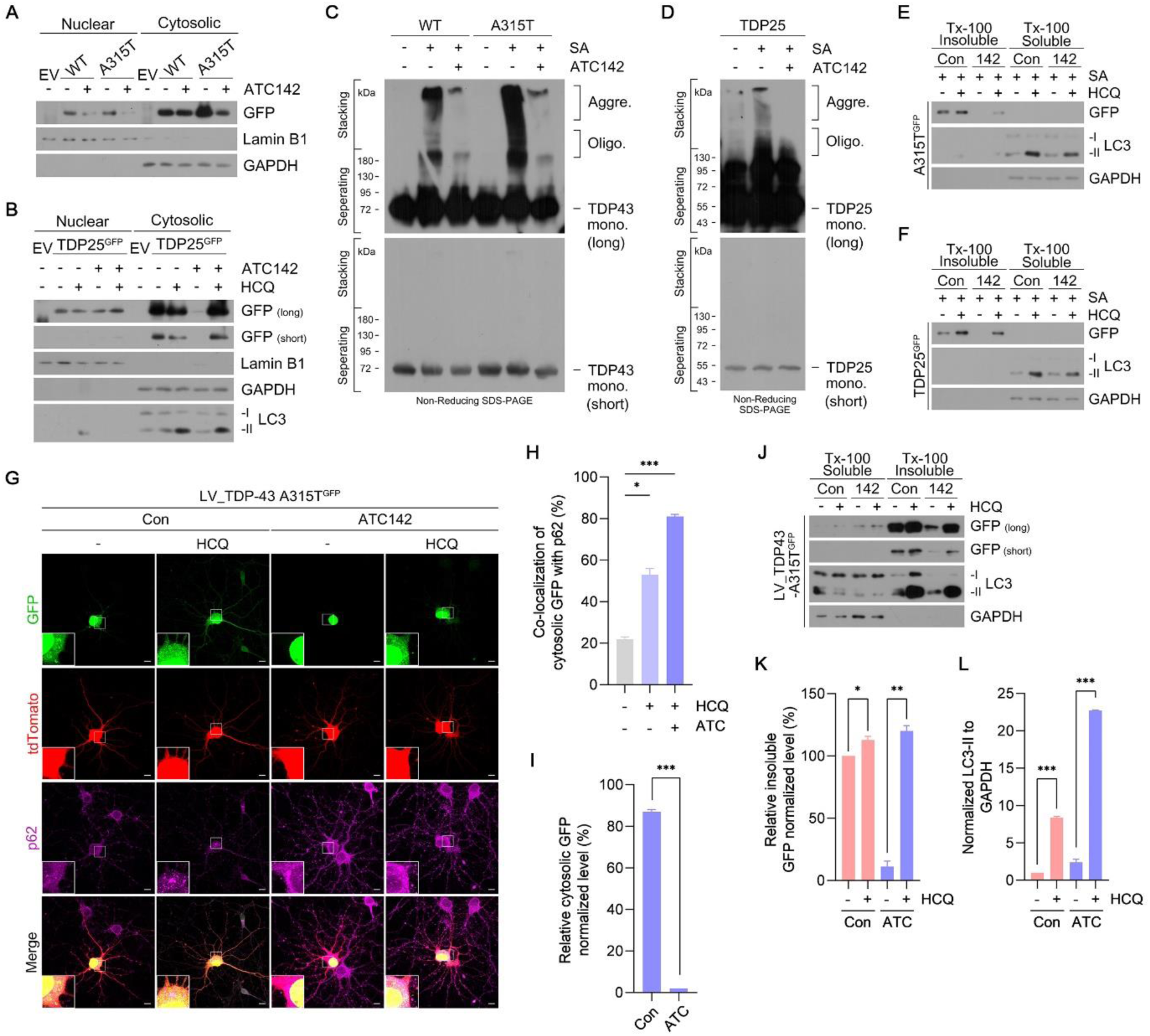
ATC142 selectively eliminates aggregated TDP-43 while preserving its functional monomers. **(A)** Nuclear/cytosolic fractionation assay in HEK293T cells transfected with TDP-43 WT-GFP or TDP-43 A315T-GFP and treated with ATC142 (1 µM, 24 h). **(B)**. Identical to (A), but in HEK293T cells transfected with TDP25-GFP and treated with or without ATC142 (1 µM, 24 h) and HCQ (10 µM, 24 h). **(C)** Non-reducing immunoblotting analysis in HEK293T cells transfected with TDP-43 WT-GFP or TDP-43 A315T-GFP, treated with ATC142 (1 µM, 24 h) and sodium arsenite (200 µM, 24 h). **(D)** Identical to (C), but in HEK293T cells transfected with TDP25-GFP. **(E)** Triton X-100 fractionation assay in HeLa cells transfected with TDP-43 A315T-GFP and treated with or without sodium arsenite (200 µM, 24 h), ATC142 (1 µM, 24 h), and HCQ (10 µM, 24 h). **(F).** Identical to (E), but in HeLa cells transfected with TDP25-GFP. **(G)** Immunocytochemistry in primary hippocampal neurons transduced with TDP-43 A315T-GFP lentivirus and treated with or without ATC142 (1 µM, 24 h) and HCQ (10 µM, 24 h). Scale bar, 10 µm. **(H)** Quantification of cytosolic co-localization between GFP and p62 in (G). **(I)** Quantification of cytosolic GFP levels in (G) (first and third columns). **(J)** Triton X-100 fractionation assay in primary cortical neurons transduced with TDP-43 A315T-GFP lentivirus and treated with or without ATC142 (1 µM, 24 h) and HCQ (10 µM, 24 h). **(K)** Quantification of GFP levels in (J). **(L)** Quantification of LC3-II levels in (J). Data are presented as mean ± S.E.M. Statistical significance is indicated as follows: *P ≤ 0.05; **P ≤ 0.01; ***P ≤ 0.001. (Student’s *t*-test).

Pathological aggregation of TDP-43 reduces its solubility in detergents, resulting in the formation of insoluble inclusion bodies (*57*). Soluble/insoluble fractionation using Triton X-100 of HEK293T cells expressing TDP-43 A315T and TDP-25 treated with sodium arsenite revealed that, under reducing conditions, only insoluble TDP-43 and TDP-25 species were detected by western blot analysis, while no soluble species were observed (Fig. 3, E and F, lanes 1-4). ATC142 treatment prevented the accumulation of insoluble TDP-43 and TDP-25 species with increased LC3 lipidation (Fig. 3, E and F). To further investigate the pathological effects of TDP-43 A315T, we generated a lentiviral vector encoding the A315T variant and delivered it into primary cultured neurons. Immunocytochemistry revealed that these neurons developed pathological TDP-43 species outside the nucleus, accumulating in both the soma and axonal compartments (fig. S4B). Subsequent imunocytochemical analyses of primary hippocampal neurons expressing TDP-43 A315T revealed co-treatment with ATC142 and HCQ increased colocalization of TDP-43 and p62 in the soma compared to HCQ alone (Fig. 3, G and H). Moreover, ATC142 significantly decreased neuritic TDP-43 aggregates, as indicated by diminished fluorescence signals (Fig. 3I). When Triton X-100-based fractionation to separate soluble and insoluble proteins in primary cortical neurons stably expressing TDP-43 A315T was performed, immunoblotting analysis showed ATC142 promoted LC3 lipidation and markedly reduced detergent-insoluble TDP-43 species by 89% under reducing SDS-PAGE conditions (Fig. 3, J to L). In conclusion, ATC142 facilitates the targeting of pathological neuritic TDP-43 species to the autophagy machinery, thereby preventing the formation of insoluble aggregates.

### ATC142 induces degradation of cytosolic TDP-43 aggregates in ALS patient-derived iPSC motor neurons

To validate the degradative efficacy of ATC142, we generated an ALS patient-derived iPSC line from skin fibroblasts. These fibroblasts were obtained from an ALS patient carrying the p.A382T pathogenic variant of the *TARDBP* gene. Fibroblasts from a healthy donor were used as the control group. Both ALS and control iPSC colonies exhibited typical round morphology characterized by small, tightly packed cells (fig. S5, A and D). Pluripotency of each line was confirmed by RT-PCR analysis, immunocytochemistry using stem cell markers (Oct4, Sox2, Nanog, Tra-1-81, Tra-1-60, or SSEA4), and alkaline phosphate activity assays (fig. S5, B to D).

To assess cytosolic TDP-43 aggregation in motor neurons derived from iPSCs, we followed a previously described differentiation protocol with minor modifications (*58*). Characterization of these motor neurons was performed by immunocytochemistry using neural differentiation markers including Sox2, Nestin, Pax6, HB9, and ChAT (fig. S6A and B). As shown in supplementary figure 6C and D, ALS-iPSC-derived motor neurons exhibited a significant accumulation of cytosolic TDP-43 aggregates compared to control neurons.

We next evaluated the degradative efficacy of ATC142 against these cytosolic TDP-43 aggregates. Immunocytochemistry revealed treatment of ALS-iPSCs with ATC142 led to a dose-dependent reduction in both the number and total area of TDP-43 aggregates by up to 2.7-fold and 2.6-fold, respectively, compared to the vehicle-treated group. Co-treatment of ATC142 with bafilomycin A1, a V-ATPase inhibitor that impairs lysosomal acidification, led to a significant increase in the number and total area of TDP-43 aggregates comparable to the control group (Fig. 4, A to C). Collectively, ATC142 promoted autophagy-dependent clearance of TDP-43 aggregates in ALS-iPSCs.

**Fig. 4.**
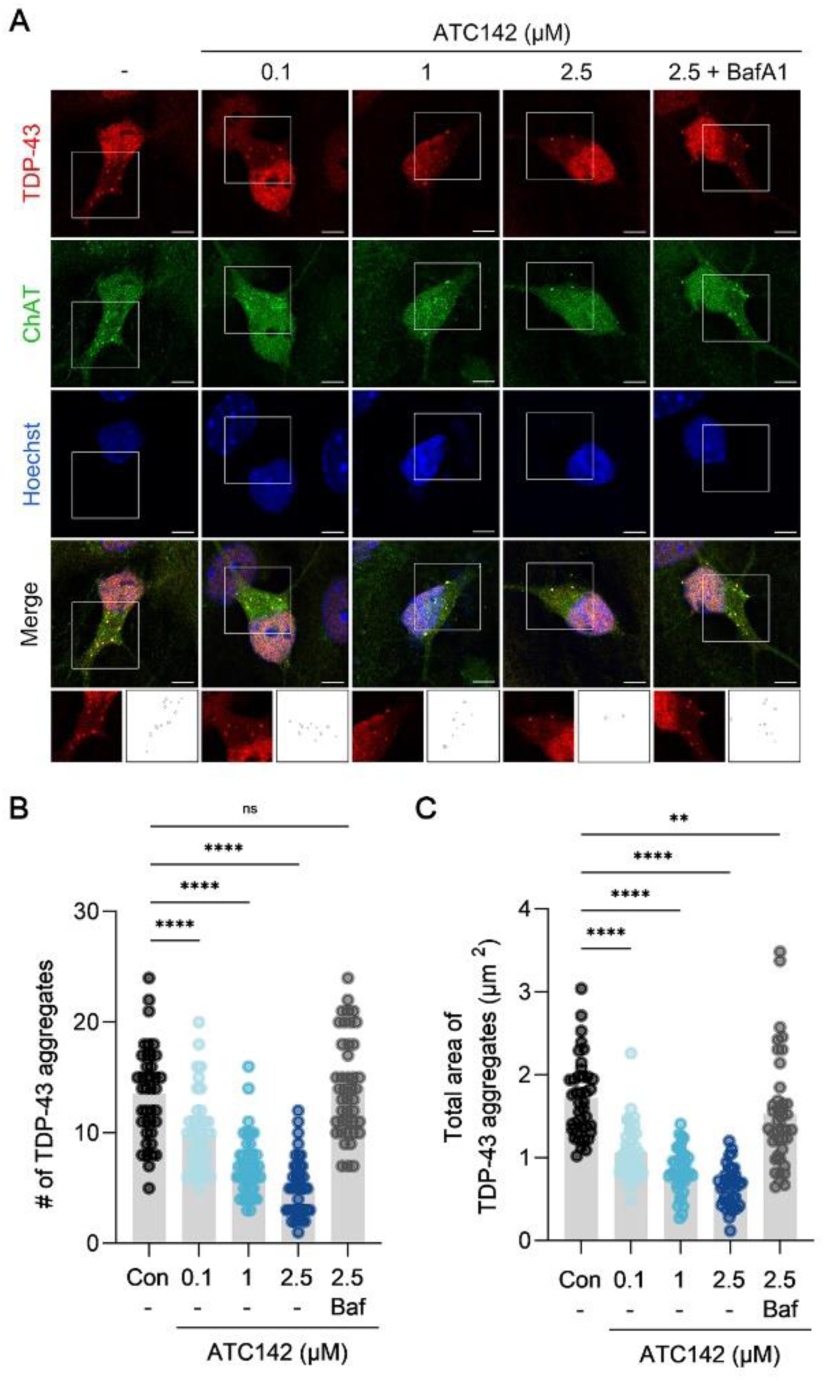
ATC142 induces degradation of cytosolic TDP-43 aggregates in ALS patient-derived iPSC motor neurons. **(A)** Immunocytochemistry of ALS iPSCs treated with ATC142 at the indicated concentrations (24 h), with or without bafilomycin A1 (10 nM, 24 h). Scale bar, 10 µm. **(B)** Quantification of the number of TDP-43 aggregates shown in (A). **(C)** Quantification of the total area of TDP-43 aggregates shown in (A). Data are presented as mean ± S.E.M. Statistical significance is indicated as follows: ns, not significant; **P ≤ 0.01; ****P ≤ 0.0001. (Student’s *t*-test).

### ATC142 suppresses TDP-43-mediated NF-κB activation

Pathological TDP-43 oligomers activate the phosphorylation of p65, a subunit of NF-κB, leading to the transcriptional activation of pro-inflammatory cytokines and the damage of motor neurons (*21, 59*). As expected, transient expression of TDP-43 A315T or TDP-25 in HEK293T cells increased the levels of phosphorylated p65 (p-p65), which were reduced by 82% and 93%, respectively, upon ATC142 treatment in a dose-dependent manner (Fig. 5, A to F). The treatment of sodium arsenite further accelerated the acute accumulation of p-p65, which was efficiently counteracted with ATC142 in a time-dependent manner (Fig. 5, G and H). Similar dose-dependence was observed with HeLa cells transiently expressing TDP-43 A315T or TDP-25 (Fig. 5, I to L). As an alternative approach, we also adopted a primary cortical neuron model transduced with TDP-43 A315T. In primary cortical neurons expressing TDP-43 A315T, sodium arsenite treatment led to a ∼7-fold increase in p-p65 levels, which was reduced by ∼72% upon ATC142 treatment (Fig. 4, M and N). These results demonstrate the efficacy of ATC142 in alleviating TDP-43-induced NF-κB activation.

**Fig. 5.**
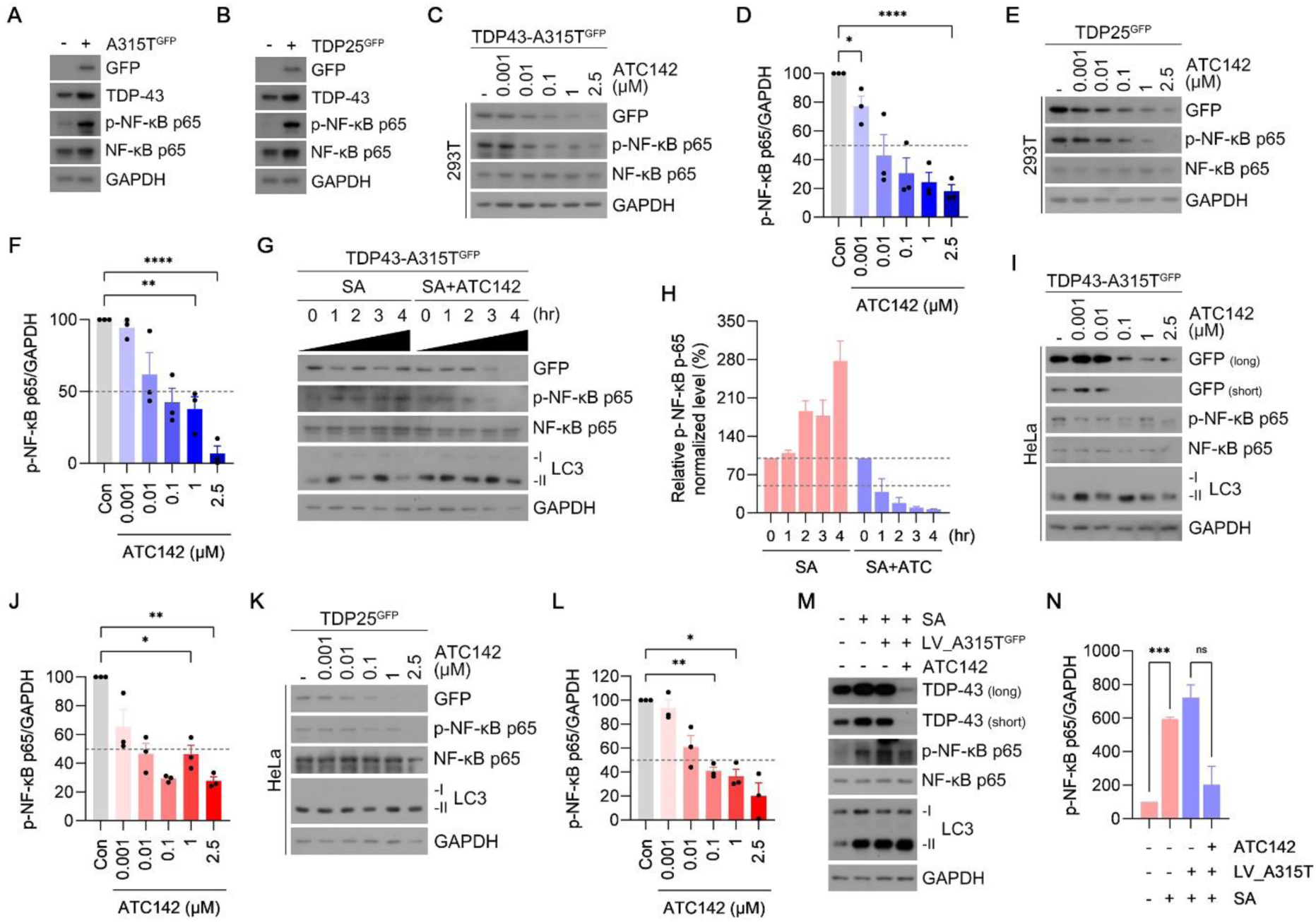
ATC142 suppresses TDP-43-mediated NF-κB activation. **(A)** Immunoblotting analysis of HEK293T cells with or without TDP-43 A315T-GFP transfection. **(B)** Identical to (A), but with or without TDP25-GFP transfection. **(C)** Immunoblotting analysis of HEK293T cells transfected with TDP-43 A315T-GFP and treated with ATC142 at the indicated concentrations (24 h). **(D)** Quantification of phospho-NF-κB p65 levels shown in (C). **(E)** Identical to (C), but in HEK293T cells transfected with TDP25-GFP. **(F)** Quantification of phospho-NF-κB p65 levels shown in (E). **(G)** Immunoblotting analysis of HEK293T cells transfected with TDP-43 A315T-GFP and treated with or without sodium arsenite (400 µM) and ATC142 (1 µM) at the indicated time points. **(H)** Quantification of phospho-NF-κB p65 levels shown in (G). **(I)** Immunoblotting analysis of HeLa cells transfected with TDP-43 A315T-GFP and treated with ATC142 at the indicated concentrations (24 h). **(J)** Quantification of phospho-NF-κB p65 levels shown in (I). **(K)** Identical to (I), but in HeLa cells transfected with TDP25-GFP. **(L)** Quantification of phospho-NF-κB p65 levels shown in (K). **(M)** Immunoblotting analysis of primary cortical neurons transduced with or without TDP-43 A315T-GFP lentivirus and treated with or without sodium arsenite (200 µM, 24 h) and ATC142 (1 µM, 24 h). **(N)** Quantification of phospho-NF-κB p65 levels shown in (M). Data are presented as mean ± S.E.M. Statistical significance is indicated as follows: ns, not significant; *P ≤ 0.05; **P ≤ 0.01; ***P ≤ 0.001; ****P ≤ 0.0001. (Student’s *t*-test).

### ATC141 induces the targeted degradation of pathogenic TDP-43 species in the spinal cord of ALS model mice

To evaluate the therapeutic efficacy of AUTOTACs in mice, we compared a set of AUTOTACs carrying Anle138b or its derivatives as the TBL for their pharmacokinetic (PK) profiles, oral availability, solubility, and toxicity in mice (table S2). This screening identified an orally administrative compound, ATC141/ATB2005A with a size of 734 Da (*60*), that was formulated into a salt form containing hydrochloride from ATB2005 with a size of 624 Da (*46, 51*). In non-reducing SDS-PAGE of HEK293T cells expressing TDP-43 A315T, ATC141 exhibited a DC_50_ of 431 nM for high molecular-weight species, but no detectible degradation activities for its monomeric form (fig. S7, A and B). When Triton X-100-based insoluble proteins were fractionated from primary cortical neurons expressing TDP-43 A315T, ATC141 induced a similar dose-dependent degradation of detergent-resistant insoluble TDP-43 species at DC_50_ of 420 nM associated with enhanced LC3 lipidation (fig. S7C). Immunocytochemistry revealed that ATC141 caused a dose-dependent reduction in the number and total area of neuritic TDP-43 aggregates by 2.6- and 3.1-fold, respectively (fig. S7, D to F). ATC141 failed to exert such an efficacy when autophagic flux was blocked using bafilomycin A1 (fig. S7, E and F, lane 5). These results show that ATC141 induces autophagic degradation of misfolded TDP-43 A315T aggregates.

Next, we assessed the degradation efficacy of ATC141 in B6.Cg-Tg(Prnp-TARDBP*A315T)95Balo/J mice expressing TDP-43 A315T in the spinal cord (*61*). These TDP-43 A315T mice develop ubiquitinated TDP-43 and TDP-25 aggregates and inclusions in spinal motor neurons, recapitulating pathological features of ALS (*61, 62*). The mice at 16 weeks of age were orally administered with 10 mg/kg ATC141 three times per week (TIW) for eight consecutive weeks, totaling 24 doses (Fig. 6A and fig. S8A). The lysates of spinal cords were separated into soluble and insoluble fractions using Triton X-100, followed by non-reducing SDS-PAGE. ATC141 administration induced the lowering of high molecular weight TDP-43 species in the insoluble fraction by ∼48% (Fig. 6, B and C). By contrast, no significant lowering was observed with monomeric TDP-43 in the soluble fraction (Fig. 6B, lanes 1-12). When the dorsal horn of spinal cord cross-sections was subjected to immunohistochemistry, ATC141 administration almost completely eliminated TDP-43-positive inclusions to a level comparable to non-transgenic control mice (Fig. 6, D and E). These results demonstrate that ATC141 effectively reduces pathological TDP-43 accumulation *in vivo*.

**Fig. 6.**
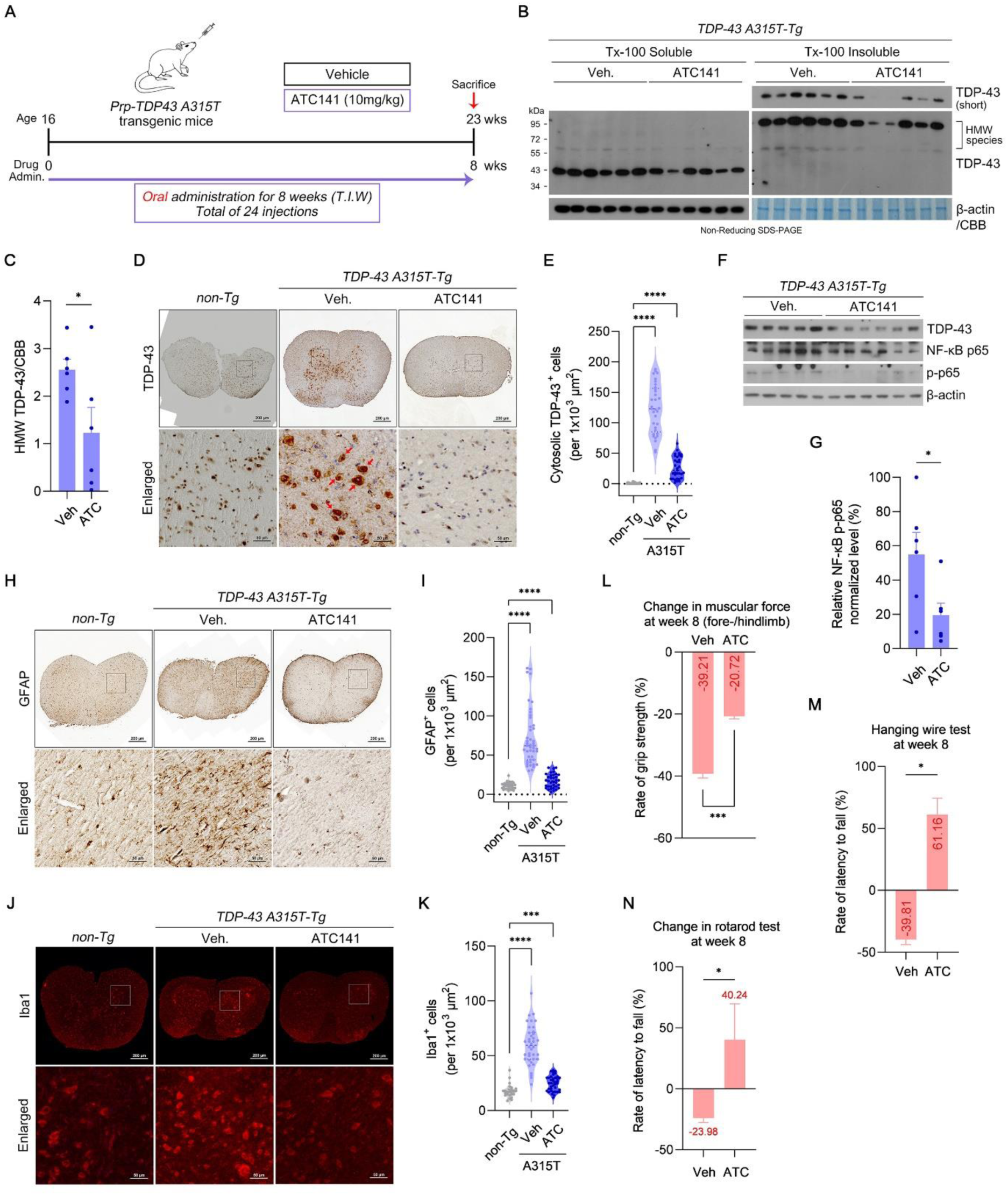
ATC141 reduces TDP-43 aggregation and alleviates TDP-43 pathology in the spinal cord of ALS model mice. **(A)** Schematic representation of the experimental timeline for Prp-TDP-43 A315T mice. Mice were administered ATC141 (10 mg/kg) starting at 16 weeks of age, three times a week for 8 weeks. Behavioral testing was performed at 16 and 23 weeks of age, followed by sacrifice for subsequent analyses. **(B)** Non-reducing SDS PAGE of Triton X-100-fractionated spinal cord lysates from vehicle- and ATC141-treated mice to detect TDP-43. **(C)** Quantification of high-molecular-weight TDP-43 in the insoluble fractions shown in (B). **(D)** Immunohistochemistry of TDP-43 in spinal cord cross-sections from non-transgenic and TDP-43 A315T mice. Scale bars: 200 µm (full view), 50 µm (enlarged). **(E)** Number of cytosolic TDP-43-positive cells per 1×10^3^ µm^2^ in (D). **(F)** Immunoblotting analysis of spinal cord lysates from TDP-43 A315T mice to detect NF-κB p65 and phospho-NF-κB p65. **(G)** Quantification of phospho-NF-κB p65 levels shown in (F). **(H)** Immunohistochemistry of GFAP in spinal cord cross-sections from non-transgenic and TDP-43 A315T mice. Scale bars: 200 µm (full view), 50 µm (enlarged). **(I)** Number of GFAP-positive cells per 1×10^3^ µm^2^ in (H). **(J)** Immunohistochemistry of Iba1 in spinal cord cross-sections from non-transgenic and TDP-43 A315T mice. Scale bars: 200 µm (full view), 50 µm (enlarged). **(K)** Number of Iba1-positive cells per 1×10^3^ µm^2^ in (J). **(L)** Comparative analysis of combined forelimb and hindlimb grip strength in TDP-43 A315T mice at week 0 and week 8. **(M)** Comparative analysis of latency to fall in the rotarod test in TDP-43 A315T mice at week 0 and week 8. **(N)** Comparative analysis of latency to fall in the hanging wire test in TDP-43 A315T mice at week 0 and week 8. Data are presented as mean ± S.E.M. Statistical significance is indicated as follows: *P ≤ 0.05; ***P ≤ 0.001; ****P ≤ 0.0001. (Student’s *t*-test).

### ATC141 exerts the therapeutic efficacy in neuroinflammation of ALS model mice expressing TDP-43 A315T

TDP-43 A315T mice develop ALS-related phenotypes in the spinal cord, characterized by glial activation and progressive motor deficits (*61, 63*). To assess the therapeutic efficacy of ATC141 in modulating neuroinflammatory responses, spinal cord lysates of ATC141-treated mice were subjected to non-reducing SDS-PAGE, followed by immunoblotting analyses. Compared with vehicle-treated mice, ATC141 treatment lowered the level of p-p65 by ∼36% (Fig. 6, F and G). To further investigate the effect of ATC141 on glial reactivity, we analyzed the expression of GFAP, an intermediate filament protein upregulated in reactive astrocytes.

Immunohistochemical analyses showed that the spinal cords of TDP-43 A315T mice contained 563% more GFAP^+^ astrocytes compared to non-transgenic controls (Fig. 6, H and I, lanes 1, 2). Following ATC141 administration, the level of GFAP^+^ astrocytes was reduced by 76% to a level comparable to those in non-transgenic mice (Fig. 6, H and I). A similar reduction was observed for Iba1, a cytoplasmic protein specifically expressed in activated microglia, as ATC141 treatment markedly reduced the number of Iba1^+^ cells by 58% (Fig. 6, J and K). Collectively, these results demonstrate that ATC141 attenuates NF-κB signaling and glial activation in the ALS mouse model.

### ATC141 ameliorates motor and muscular deficits in TDP-43 A315T mice

To validate the therapeutic efficacy of ATC141 in neuromuscular functions, TDP-43 A315T mice at 16 weeks of age were orally administered with 10 mg/kg ATC141 TIW for 8 weeks, followed by a series of behavioral tests. At this age, TDP-43 A315T mice exhibited motor deficits as evidenced by a 31% decrease in forelimb grip strength and a 50% reduction in rotarod latency as compared to non-transgenic controls (fig. S8, B and C). After 8-weeks of administration, whereas forelimb grip strength of vehicle-treated mice decreased by 41%, from 88.8 g to 52.7 g, ATC141-treated mice lost only 29%, from 88.9 g to 63.2 g (fig. S8, D and E). A direct comparison of post-treatment trajectories between the two groups revealed 28.8% improvement (fig. S8E). When the grip strengths of forelimbs and hindlimbs were combined, ATC141-treated mice exhibited a 21% reduction (179.0 g to 141.9 g) as compared with a 39% reduction (172.5 g to 104.8 g) of vehicle-treated mice (Fig. 6L and fig. S8F), indicating therapeutic efficacy of 45.2% when compared between the two groups. These results demonstrate the therapeutic efficacy of ATC141 in muscle strength.

To assess the efficacy of ATC141 in neuromuscular coordination, we conducted rotarod tests using an accelerating protocol (4 to 20 rpm over 120 s). Vehicle-treated mice showed a 24% decline in latency to fall over 8 weeks of administration (Fig. 6M and S8G). Strikingly, ATC141-treated mice exhibited a 40% increase under the same conditions, representing a 268% relative improvement compared to vehicle-treated controls. These results indicate that ATC141 can reverse the disease progression of ALS. To further assess motor performance, we conducted a hanging wire test in which mice grasped the center of a horizontal wire with both forelimbs. When the latency to fall was recorded over 120 seconds, vehicle-treated mice showed a 40% decrease over 8 weeks of treatment (Fig. 6N and S8H). ATC141-treated mice exhibited a therapeutic improvement of 61% as compared with Day 1, corresponding to a 254% relative improvement compared to vehicle treatment. As an alternative measurement, mice were scored on a 0-4 success scale based on their performance and final position on the wire. The mean success score of ATC141-treated mice improved markedly from 1.11 to 3.25, while that of vehicle-treated mice declined from 1.67 to 1.27 over the same period. This corresponds to a 635% relative improvement in mean success score for ATC141-treated mice compared to controls (fig. S8, I to K). Finally, we monitored the impact of ATC141 administration on the life-span of the mice. ATC141-treated mice exhibited the median survival of 191 ± 9 days as compared with 174 ± 12 days of the vehicle group (fig. S8L). These results show that ATC141 not only halt the disease progression but also at least partially cure the disease and prolongs survival of TDP-43 A315T mice.

## Discussion

The ALS is a fatal progressive disease that results in the degeneration of motor neurons in the spinal cord (*1*). Despite desperate studies over past decades to at least delay the disease progression, there are no effective drugs available in the market (*4*). Although ALS is initially caused by heterogenous factors such as oxidative stress and mitochondrial dysfunction in sALS and genetic mutations in fALS, almost all the ALS cases end up with the accumulation of misfolded protein aggregates (*64*). In this study, we employed AUTOTAC to degrade pathogenic aggregates of TDP-43 species founded in the spinal cord of 97% of all ALS cases. ATC141 and ATC142 were designed to simultaneous bind β-sheet-rich TDP-43 aggregates and the autophagic N-recognin p62 leading to autophagosomal sequestration and lysosomal co-degradation (Fig. 1, A and B, and Fig. 2, C and D). Both compounds exhibited degradative efficacy against the pathogenic species of TDP-43 A315T and TDP-25 with DC_50_ values ranging from low nM to sub-μM depending on physiological and experimental conditions (Fig. 1, C to F, and fig. S3, A to D). In ALS model mice expressing TDP-43 A315T in the spinal cord, oral administration of 10 mg/kg ATC141 with 24 doses were sufficient to lower pathological TDP-43 species and neuroinflammation, which correlated to the therapeutic improvements in motor and muscular deficits (Fig. 6, C to M, and fig. S8, A to E). Importantly, ATC141 reversed the disease progression and prolonged the survival of TDP-43 A315T mice (Fig. 6N and O, and fig. S8, F to K). Our results suggest that AUTOTAC provides a novel therapeutic strategy to develop drugs for a broad spectrum of proteinopathies including neurodegenerative diseases.

The pathogenesis of ALS is initially driven by various mechanisms ranging from neuroinflammation and oxidative stress to autoimmunity and mitochondrial dysfunction, which eventually leads to the misfolding and aggregation of ALS-linked proteins (*14, 16*). However, despite extensive efforts to develop therapeutic means that intervene with these ALS-associated pathophysiological processes, currently available drugs are at best marginally effective to delay the disease progression (*23, 24*). As an alternative approach, recent studies employed TPD technologies as represented by PROTAC (*27*). Indeed, a PROTAC designed to target TDP-43 aggregates was shown to improve the motility in a *Caenorhabditis elegans* ALS model (*28*), suggesting its potential to degrade misfolded TDP-43 monomers as a preventive medicine. However, TDP-43 aggregates are structurally stable and resistant to unfolding, making them poor substrates for proteasomal degradation (*13, 15*). In addition, the narrow pore size of the proteasome further limits its ability to clear these aggregates (*31*). To overcome these limitations, we developed AUTOTAC, which leverages p62-mediated macroautophagy for lysosomal degradation (*46*). In contrast to PROTAC that strictly depends on the formation of an E3-substrate tertiary complex, an AUTOTAC compound acts as an agonist that induces the allosteric conformational change of p62 independent of its engagement with cargoes (*27*). As p62 undergoes self-polymerization via the liquid-liquid phase separation (LLPS), its cargoes are co-polymerized through multivalent interactions. The p62-cargo complexes are transiently sequestrated by phagophores and autophagosomes, followed by the fusion between autophagosome and lysosomes, leading to co-degradation by lysosomal hydrolases (*46*). Despite these multi-step mechanisms underlying AUTOTAC and the prevailing notion that autophagy is a bulky, slow, and nonspecific process, ATC142 induced lysosomal degradation of pathogenic TDP-43 species by over 90% with DC_50_ of 1.0-25 nM in an acute manner (Fig. 1, C to F, I to L, and fig. S3, A to D). Such an acute, potent degradation activity can be attributed to the activity of p62-mediated macroautophagy to degrade large complexes at once by lysosomal hydrolases. We suggest that autophagy provides an ultimate solution to eliminate pathogenic agents in protein and cellular wastes.

The disappointingly modest effects of FDA-approved riluzole and edaravone, which act to rectify excitotoxicity and oxidative stress, imply a more central pathway needs to be targeted (*23, 24, 65*). Consequently, over the past decades, extensive studies explored autophagy to degrade ALS-related protein aggregates including FUS, SOD1, TDP-43, and dipeptide repeats from *C9ORF72* (*26, 33, 34*). Rapamycin, an inhibitor of the mammalian target of rapamycin complex 1 (mTORC1), is the most widely recognized pharmacological autophagy inducer and was first reported in 2011 to induce autophagy in SOD1 mutant mice (*66*). However, while rapamycin rescued motor neuron function in mice overexpressing wild-type TDP-43, it exhibited detrimental effects in SOD1 transgenic models (*66–68*). These findings suggest that global mTORC1 inhibition may not be broadly beneficial for ALS, especially given that mTORC1 is a master regulator of protein synthesis, immune responses, and cell cycle progression, and prolonged treatment can inhibit mTORC2 (*69*). To overcome these limitations, mTOR-independent autophagy inducers, including trehalose (*70, 71*) and rilmenidine (*72*), have been evaluated in SOD1 mutant mice, but yielded inconsistent outcomes, indicating that bypassing mTOR signaling to induce autophagy remains challenging. Notably, rilmenidine induced autophagy in the spinal cord but paradoxically exacerbated paralysis and reduced lifespan in a TDP-43 transgenic mouse model (*73*). Autophagy inducers were shown to demonstrate variable efficacy depending on the disease model, act at different stages of the autophagy pathway, and often affect multiple biological processes, which may result in off-target toxicity (*22, 34, 74*). In this study, we assessed the therapeutic efficacy of ATC141 *in vivo* by employing transgenic mice expressing aggregation-prone human TDP-43 A315T. This model uses the PrP promoter to drive TDP-43 A315T expression in both motor neurons and glial cells, thereby recapitulating cell-autonomous and non-cell autonomous mechanisms implicated in ALS. These include progressive accumulation of ubiquitinated TDP-43 and TDP-25 in the spinal cord, motor neuron and axonal degeneration, astrogliosis, and microglial activation, all contributing to motor dysfunction (*61–63*). Our results show that total 24 oral doses of 10 mg/kg ATC141 TIW were sufficient to lower detergent-insoluble TDP-43 aggregates in the spinal cord (Fig. 6, C and D) and attenuated neuroinflammation as evidenced by astrogliosis and microglial activation (Fig. 6, E to L). These molecular improvements translated into functional benefits, with a 268% relative improvement in rotarod performance and a 635% enhancement in hanging wire test outcomes (Fig. 6, N and O, and fig. S8, F to J). Importantly, this oral drug was able to not only delay or halt the diseases progression but at least partially cure the disease. These results provide a possibility that motor neurons denervated from muscles in ALS patients can restore their neuromuscular functions if pathogenic protein aggregates are properly eliminated. Moreover, further therapeutic efficacy could be expected if the drug is administered at higher doses and for a longer period of time. Through cross seeding and co-aggregation of ALS proteins, TDP-43 proteinopathies underly in other neurodegenerative disorders, including AD (*20*), PD (*17*), and FTD (*21*). Thus, ATC141 may be applied for a broad spectrum of neurodegenerative diseases involving TDP-43 aggregation. Originally designed to recognize β-sheet rich oligomers, ATC141 is under Phase 1 clinical trials in South Korea with 76 healthy volunteers (IND No. 2023-00697) in preparation of Phase 2 trials to treat a broad spectrum of proteinopathies such as AD, PSP, and ALS. As a veterinary medicine, Phase 2 trials demonstrated the therapeutic efficacy of ATC141 to reverse the disease progression of dog patients carrying canine cognitive dysfunction (CCD), an equivalent to human AD (U34401-4/2023/14). Given that β-sheet rich oligomers are commonly found in many neurodegenerative diseases, ATC141 and related AUTOTAC compounds may provide a therapeutic means for a broad spectrum of currently incurable diseases involving protein aggregates.

Despite the compelling preclinical efficacy of ATC141 and ATC142 in reducing pathological TDP-43 aggregates and ameliorating disease phenotypes, several considerations remain for clinical translation. The long-term safety and potential off-target effects of AUTOTAC compounds require thorough evaluation, particularly given the critical physiological roles of p62 and autophagy. Interestingly, we observed reaccumulation of GFP-tagged TDP-43 signal following AUTOTAC washout. This raises several mechanistic questions, including whether the resurgence reflects ongoing synthesis of TDP-43 or a limitation in the catalytic recycling of the AUTOTAC molecule. Alternatively, it may indicate a threshold in autophagic flux capacity, suggesting that repeated dosing could be necessary to sustain therapeutic benefit in clinical settings. Although ATC141 and ATC142 share the same TBL, they exhibited variable degradative efficacy, likely due to differences in ATL positioning and compound permeability, both of which influence p62 recruitment and autophagic flux. This underscores the importance of rational design and optimization of AUTOTAC compounds to improve p62 engagement and autophagic flux. Also, while AUTOTAC efficacy was validated in patient-derived iPSC motor neurons and in an in vivo ALS model, many of our cellular assays utilized plasmid-based overexpression to induce cytoplasmic TDP-43 accumulation. Such artificial models may not fully recapitulate the complexity of endogenous aggregation dynamics in ALS patients. Future studies should address these challenges to enable broader clinical application. Nonetheless, our results provide a compelling proof-of-concept that p62-mediated selective macroautophagy establishes a foundation for the development of future targeted degradation therapies, addressing a critical unmet need in ALS and other TDP-43 proteinopathies.

## Materials and Methods

### Cell culture

HEK293T and HeLa cells were obtained from the Korean Cell Line Bank. Cells were maintained in Dulbecco’s Modified Eagle’s Medium (DMEM; 11995073, Gibco) supplemented with 10% fetal bovine serum (FBS; 16000044, Gibco) and antibiotics (100 units/mL penicillin-streptomycin). Cells were incubated at 37° in a 5% CO_2_ humidified chamber.

### Antibodies

The following primary antibodies were used: mouse monoclonal anti-SQSTM1/p62 (1:10,000 for WB, 1:250 for ICC; ab56416, Abcam), rabbit polyclonal anti-LC3 (1:10,000 for WB; 1:150 for ICC; L7532, Sigma-Aldrich), rabbit polyclonal anti-GAPDH (1:10,000 for WB; AP0066, BioWorld), rabbit polyclonal anti-β-actin (1:10,000 for WB; AP0060, BioWorld), mouse monoclonal anti-Lamin B1 (1:5,000 for WB; sc-374015, Santa Cruz), mouse monoclonal anti-ChAT (1:200 for ICC; MA5-31383, Invitrogen), rabbit monoclonal anti-TDP-43 (1:5,000 for WB, 1:200 for IHC; ab109535, Abcam), rabbit polyclonal anti-ATG5 (1:1,000 for WB; NB110-53818, Novus), rabbit monoclonal anti-ATG7 (1:1,000 for WB; 8558, Cell Signaling Technology), mouse monoclonal anti-Ub (1:2,000 for WB; sc-8017, Santa Cruz), mouse monoclonal anti-GFP (1:1,000 for WB; sc-9996, Santa Cruz), rabbit polyclonal anti-Myc (1:5,000 for WB; 2278, Cell Signaling Technology), rabbit polyclonal anti-Flag (1:5,000 for WB; F7425, Sigma-Aldrich), rabbit monoclonal anti-NF-κB p65 (1:5,000 for WB; 4764, Cell Signaling Technology), rabbit monoclonal anti-phospho NF-κB p65 (1:5,000 for WB; 3033, Cell Signaling Technology), rabbit monoclonal anti-GFAP (1:200 for IHC; A19058, Abclonal), and rabbit polyclonal anti-Iba1 (1:200 for IHC; 019-19741, Wako). The following secondary antibodies were used: alexa fluor 488 goat anti-rabbit IgG (1:1,000; A11034, Invitrogen), alexa fluor 488 goat anti-mouse IgG (1:1,000; A11029, Invitrogen), alexa fluor 555 goat anti-rabbit IgG (1:1,000; A21429, Invitrogen), alexa fluor 555 goat anti-mouse IgG (1:1,000; A32727, Invitrogen), alexa fluor 633 goat anti-mouse IgG (1:1,000; A21052, Invitrogen), anti-rabbit IgG-HRP (1:10,000, 7074, Cell Signaling Technology), and anti-mouse IgG-HRP (1:10,000; 7076, Cell Signaling Technology).

### Primary neuron culture

Primary cortical or hippocampal neurons were isolated from embryonic day 18 (E18) Sprague Dawley rats (OrientBio). All procedures involving animals were approved by the Seoul National University Institutional Animal Care and Use Committee (protocol No. SNU-161222-2-4) and conducted in accordance with institutional guidelines. Rats were house individually in standard cages with *ad libitum* access to food and water, and maintained under controlled conditions (22 ± 2°, 50 ± 10% humidity, 12-h light/dark cycle). Pregnant rats were sacrificed by CO_2_ asphyxiation followed by decapitation to minimize suffering. Embryos were delivered via Caesarean section and decapitated. The hippocampi or cortices were dissected and incubated in Hanks’ Balanced Salt Solution (HBSS; 14170-161, Thermo Fisher Scientific) containing 0.05% trypsin (T1005, Sigma-Aldrich), 10 mM HEPES (H3375, Sigma-Aldrich), 0.137 mg/mL DNase I (D5025, Sigma-Aldrich), and penicillin-streptomycin (15070-063, Thermo Fisher Scientific) for 12 min at 37°. Tissues were gently dissociated by trituration using a fire-polished Pasteur pipette and plated on poly-D-lysine-coated culture plates in serum-free Neurobasal medium (21103-049, Thermo Fisher Scientific) supplemented with B-27 (17504-044, Thermo Fisher Scientific) and 1% L-glutamine (G7513, Sigma-Aldrich). Neurons were maintained at 37° in a 5% CO_2_ humidified incubator, and the medium was partially replaced with fresh medium every 2-3 days.

### Plasmid DNA and cloning

Plasmids harboring human TDP-43 wild-type (WT), TDP-43 A315T mutant, and TDP-25 (a C-terminal fragment of TDP-43) were obtained as gifts from Jin-A Lee (Hannam University), and subcloned into the pEGFP-C3 vector to generate TDP-43 WT-EGFP, TDP-43 A315T-EGFP, and TDP-25-EGFP. For stable overexpression of exogenous TDP-43 variants, TDP-43 WT-EGFP and TDP-43 A315T-EGFP constructs were further subcloned under the human ubiquitin C (UbC) promoter into the FUW lentiviral vector. Lentiviral particles were produced in HEK293T cells via co-transfection of the FUW constructs with the packaging plasmid △8.9 and the envelope plasmid VSV-G. Supernatants containing viral particles were collected 60 h after transfection and concentrated by ultracentrifugation at 120,000 x g for 15 min at 4°. The viral pellet was re-suspended in PBS. All DNA constructs were verified by Sanger sequencing.

### Transfection

HEK293T and HeLa cells were transfected with plasmid DNA using Lipofectamine 2000 Transfection Reagent (Invitrogen) following the manufacturer’s protocol. Predesigned siRNAs, including siRNAs (sip62, siATG5, siATG7, si*Ubb*) from Bioneer, were transfected into HeLa cells using RNAiMAX Transfection Reagent (13778075, Invitrogen) following the manufacturer’s instructions.

### Immunoblotting

Cells or tissue pellets were lysed in radioimmunoprecipitation assay (RIPA) buffer (50 mM Tris-HCl, pH 8.0, 150 mM NaCl, 1% NP-40, 0.5% sodium deoxycholate, and 0.1% sodium dodecyl sulfate (SDS)) supplemented with protease inhibitor cocktails (Sigma-Aldrich). Lysates were centrifuged at 13,000 x g for 10 min at 4°, and protein concentrations were determined using the Pierce BCA Protein Assay Kit (Thermo Fisher Scientific). For reducing SDS-PAGE, protein samples were mixed with 4 x Laemmli sample buffer (Bio-Rad) containing β-mercaptoethanol and boiled at 110° for 7 min. For non-reducing SDS-PAGE, protein lysates were mixed with non-reducing 4x LDS sample buffer (277.8 mM Tris-HCl, pH 6.8, 4.4% LDS, 44.4% (v/v) glycerol) without reducing agents and boiled at 100° for 10 min. Samples were loaded onto a 3% stacking and 8% separating SDS-PAGE gel. Proteins were separated by SDS-PAGE and transferred to polyvinylidene difluoride (PVDF) membranes at 100 V for 2 h at 4°. Membranes were blocked with 4% skim milk in PBST (PBS with 0.1% Tween-20) for 30 min at room temperature and incubated overnight at 4° with primary antibodies. After washing, membranes were incubated with HRP-conjugated secondary antibodies and developed using enhanced chemiluminescence (ECL) reagents and X-ray film. Band intensities were quantified using ImageJ software (NIH).

### Immunocytochemistry (ICC)

Cells were cultured on poly-L-lysine-coated coverslips (P4832, Sigma-Aldrich) for immunofluorescence analysis. Cells were fixed with 4% paraformaldehyde in PBS (pH 7.4) for 15 min at room temperature and washed three times with PBS for 5 min each. After fixation, cells were permeabilized with 0.5% Triton X-100 in PBS for 15 min and washed as above. Non-specific binding was blocked by incubating cells in 2% BSA/PBS for 1 h at room temperature. Cells were then incubated with primary antibodies overnight at 4°. After washing three times with PBS for 10 min each, cells were incubated with Alexa Fluor-conjugated secondary antibodies diluted in 2% BSA/PBS for 30 min at room temperature. Nuclei were counterstained with DAPI-containing mounting medium (H-1200, Vector Laboratories), and coverslips were mounted on glass slides. Confocal images were acquired using a LSM 510 Meta laser scanning confocal microscope (Zeiss) and analyzed with the Zeiss LSM Image Browser software (v4.2.0.121).

### Co-immunoprecipiation

Cells were transfected with the indicated constructs using Lipofectamine 2000 (Invitrogen) for exogenous co-immunoprecipitation (co-IP). After 24 h, cells were treated with the specified reagents for the indicated times for both endogenous and exogenous co-IP. Cells were scraped, pelleted by centrifugation, and lysed in immunoprecipitation (IP) buffer [50 mM Tris-HCl (pH 7.5), 150 mM NaCl, 0.5% Triton X-100, 1 mM EDTA, 1mM PMSF (Roche), and protease inhibitor cocktail (Sigma)] by rotation at 4° for 30 min. Lysates were sheared by passing through a 26-gauge 1 mL syringe ten times and centrifuged at 13,000 x g for 10 min at 4°. The resulting supernatants were precleared by incubation with normal mouse IgG and protein A/G-Plus agarose beads (sc-2003, Santa Cruz) overnight at 4°. For immunoprecipitation, cleared lysates were incubated with either M2 FLAG-affinity agarose beads (A2220, Sigma-Aldrich) or Myc magnetic beads (88842, Thermo Fisher Scientific) for 3 h at 4° with rotation. Beads were then washed four times with IP buffer, and bound proteins were eluted in 2x Laemmli sample buffer by boiling at 100° for 5 min. Samples were analyzed by SDS-PAGE followed by immunoblotting.

### Triton X-100 soluble/insoluble fractionation

Cells were collected using a cell lysis buffer (20 mM HEPES pH 7.9, 0.2 M KCl, 1 mM MgCl_2_, 1 mM EGTA, 1% Triton x-100, 1% SDS, 10% glycerol, protease inhibitors and phosphatase inhibitors) and incubated on ice for 15 minutes, followed by centrifugation. The supernatant was collected as the soluble fraction and the pellet as the insoluble fraction. The insoluble fraction was washed four times with PBS and then lysed with a SDS-detergent lysis buffer (20 mM HEPES pH 7.9, 0.2 M KCl, 1 mM MgCl_2_, 1 mM EGTA, 1% Triton x-100, 1% SDS, 10% glycerol, protease inhibitors and phosphatase inhibitors). Both the soluble and insoluble samples were mixed with 5X Laemmli sample buffer, boiled, and loaded onto a SDS-PAGE gel.

### Nucleus/cytoplasm fractionation

HEK293T cells were harvested, washed in PBS, and lysed in ice-cold hypotonic buffer (10 mM HEPES, 1.5 mM MgCl_2_, 10 mM KCl, 0.5 mM DTT, 0.05% NP40, protease inhibitor cocktail [Sigma-Aldrich], and 1mM PMSF [Roche]; pH 7.9) for 10 min. After centrifugation at 1,000 x g for 10 min at 4°, the supernatant (cytoplasmic fraction) was collected and mixed with 5X Laemmli sample buffer. The pellet (nuclear fraction) was resuspended in ice-cold hypertonic buffer (5 mM HEPES, 1.5 mM MgCl_2_, 4.6 mM NaCl, 0.2 mM EDTA, 0.5 mM DTT, 26% (v/v) glycerol, protease inhibitor cocktail [Sigma-Aldrich], and 1 mM PMSF [Roche]; pH 7.9), and homogenized using a glass homogenizer with 20 full strokes. The homogenate was incubated on ice for 30 min and centrifuged at 24,000 x g for 20 min at 4°. The supernatant (nuclear fraction) was collected and lysed in 5X Laemmli sample buffer.

### Immunohistochemistry

Paraffin-embedded sections of spinal cord tissue were deparaffinized in xylene and rehydrated in descent alcohol series. Antigen retrieval was performed at 95°C for 15 min using citrate buffer (pH = 6.8) and endogenous peroxidase activity was quenched in 3% hydrogen peroxide in methanol. Sections were blocked using 10% fetal bovine serum, 1% bovine serum albumin, and 0.5% Triton X-100 in PBS. After washing with PBS, sections were incubated with primary antibodies (TDP-43, GFAP, Iba1) overnight at 4°, followed by secondary anti-rabbit IgG and secondary Goat anti-rabbit AlexaFluor 488 for 1 h at room temperature. Signal detection and nucleus counter-staining were performed using the DAB substrate kit (SK-4105, Vector Laboratories) and hematoxylin QS (H-3404-100, Vector Laboratories), respectively. The stained slides were then covered with cover glass using mounting solution (H-5000-60, Vector Laboratories). Images were processed by Axio Scan Z1 (Carl Zeiss).

### iPSC generation from control and ALS patient derived primary fibroblast

iPSCs were generated from control and ALS patient-derived primary fibroblasts. The fibroblasts, obtained from the Coriell Institute for Medical Research (NINDS collection), were reprogrammed using a non-integrating plasmid induction method with OSKM factors (Healthy control: ND29178, ALS patient: ND41003). The fibroblasts were cultured at 37 °C in Dulbecco’s modified Eagle’s medium (10-013-CV, Corning) supplemented with 15% fetal bovine serum (35-015-CV, Corning) and 1% penicillin–streptomycin (SV30010, Hyclone). For gene, delivery, electroporation (Invitrogen™ Neon™ Transfection System) was employed. Seven days after electroporation, the cells were transferred onto mouse embryonic fibroblast (MEF) feeder cells with iPSC maintenance medium containing 20% knockout serum (10828028, Gibco), β-mercaptoethanol (21985023, Gibco), Gluta-MAX (35050-061, Gibco), MEM-Non Essential Amino Acids solution (11140050, Gibco), penicillin/streptomycin (SV30010, Hyclone), and DMEM-F12 (11320033, Gibco). After 3 weeks, iPSC-like colonies were selected and cultured on new feeder cells. At iPSC passage numbers 5-7, the culture system was transitioned to a feeder-free system using a plate coated with truncated recombinant human vitronectin (A14700, Gibco) and Essential 8 Medium (A1517001, Gibco).

### Characterization of the iPSCs

For immunostaining, cells were washed with DPBS, fixed with 4% PFA for 15 minutes, and permeabilized with 0.1% Triton X-100 for 15 minutes. Subsequently, they were blocked with 5% normal goat serum for 1 hour at room temperature before overnight incubation with antibodies at 4°C. Following this, cells were incubated with fluorescent-conjugated secondary antibodies (Jackson ImmunoResearch Laboratories) for 2 hours at room temperature. Finally, the cells on the glass slides were washed twice with DPBS, and the slides were mounted with mounting medium (H-1200, Vectashield). The preparations were analyzed using confocal microscopy (ZEISS, LSM 900). Antibody information: Stem cell markers: Oct3/4 (1:50; sc-5279, Santa Cruz), Nanog (1:100; RCAB003P-F, Reprocell), TRA-1-60 (1:100; MAB4360, Millipore), TRA-1-81 (1:100; MAB4381, Millipore), Neuronal markers: SOX2 (1:200; ab97959, Abcam), PAX6 (1:200; 561664, BD Pharmingen), ChAT (1:200; AB144P, Millipore), HB9 (1:6; 25179941, DSHB), Anti-TDP-43 (1:200; 3448, Cell signaling). For RNA extraction and qRT-PCR, mRNA was extracted from iPSCs using the total RNA Extraction Miniprep Kit (T2010S, NEB), and cDNA synthesis was carried out using the LunaScript RT Supermix Kit (E3010L, NEB). Real-time quantitative PCR was then performed with the QuantStudio 1 Real-Time PCR System (Thermo Fisher Scientific). Pluripotency marker expression was assessed using TaqMan probes (SOX2, Hs00602736_s1; Nanog, Hs02387400_g1; Oct3/4, Hs04260367_Gh). Regarding alkaline phosphatase analysis, alkaline phosphatase (AP) was detected using the AP Staining Kit (AP100R-1, System Biosciences) following the manufacturer’s instructions. Stemness was confirmed through alkaline phosphatase (AP) staining.

### Motor neuron differentiation

To generate highly pure motor neurons, a modified method based on Zhong-Wei Du et al., 2015, was employed. iPSCs were initially dissociated with Accutase (AT104, Innovative Cell Technologies) and split 1:6 on a Matrigel-coated plate to differentiate into neuronal precursor cells (NPC). On the following day, the Essential 8 medium was replaced with a chemically defined neural induction medium (NIP medium), consisting of Knockout DMEM-F12 (12660-012, Gibco), Neurobasal medium (21103-049, Gibco) at a 1:1 ratio, supplemented with 0.5x N2 supplement (17502-048, Gibco), 0.5x B-27 supplement (17504-044, Gibco), 1x GlutaMax supplement (35050-061, Gibco), 1x penicillin–streptomycin (SV30010, Hyclone), 100µM Ascorbic acid (A4403, Sigma), 3µM CHIR-99021 (72054, STEMCELL Technologies), 2µM DMH-1 (73634, STEMCELL Technologies), and 2µM SB-4315429 (72234, STEMCELL Technologies). The culture medium was changed every other day for 6 days. To induce ventral neurons, at 7 days after NPC cells were dissociated with Accutase and split 1:3 on a Matrigel-coated plate with NIP medium added 0.1µM Retinoic acid (R2625, Sigma) and 0.5µM Purmorphamine (72204, STEMCELL Technologies). The culture medium was changed every other day for 7 days. To induce caudal motor neurons, at 14 days after cells were dissociated with Accutase, they were cultured in suspension in NIP medium added 0.5µM Retinoic acid and 0.1µM Purmorphamine. The culture medium was half changed every other day for 7 days. To mature the motor neurons, cell clumps were dissociated with Accutase into single cells and plated on mouse glial cells with motor neuron mature medium. This medium included Neurobasal medium, 1x B-27 supplement, 1x GlutaMax, 1x penicillin–streptomycin (SV30010, Hyclone), 100µM L-Ascorbic acid (A92902, Sigma), 2.6µg/ml Adenosine 3’,5’-cyclic monophosphate (A9501, Sigma), 0.5µM Compound E (73954, STEMCELL Technologies), 10µg/ml Laminin (L2020, Sigma), 10µg/ml BDNF (450-02-50UG, Thermo Fisher Scientific), and 10µg/ml GDNF (PHC7041, Thermo Fisher Scientific). Once a week, the motor neuron mature medium was added, and the cells were matured for 5 weeks.

### Animal breeding and care

Experimental procedures were approved by the Institutional Animal Care and Use Committee (IACUC) of Seoul National University (SNU-200923-1-2). All animals received humane care and were housed in a specific-pathogen-free barrier facility at Seoul National University according to the criteria outlined in the Guide for the Care and Use of Laboratory Animals. Transgenic mice with the A315T human TDP-43 mutation under the control of the mouse prion promoter and an N-terminal Flag tag were originally developed by Dr. Robert Baloh (Washington University School of Medicine, St. Louis, MO). TDP43 A315T mutant mice were purchased at The Jackson Laboratory (stock number 010700) and extracted ear DNA amplified by the TDP43 A315T gene specific primers confirmed the TDP43 A315T carriers.

### Behavioral test

The behavioral tests were performed according to IACUC approval of Seoul National University (SNU-200923-1-2). For the grip strength test, a grip strength meter (BIO-GS3, Bioseb) was used. Three consecutive measurements were done for each mouse, and the average was calculated. For the motor coordination test, a rotarod (BX-ROD, Bioseb) was used. Each trial consisted of three attempts. The rotating speed was gradually increased from 4 to 40 rpm in the total time of 120 s. For the hanging wire test, each mouse was placed at the center of a standard metal coat hanger (2 mm in diameter, 40 cm in length) elevated approximately 40 cm above a padded surface. Mice were allowed to grasp the wire with their forelimbs and were observed for a maximum of 120 s. Motor performance was scored on a 0-4-point scale, where higher scores reflected better performance: 0 (fall within 3 s), 1 (hanging by fore- and hindlimbs >3 s), 2 (reached end of bar), 3 (climbed on top), and 4 (climbed onto angled hook).

### In silico modeling and molecular docking

The crystal structure of human TDP-43 (PDB ID: 7PY2) was retrieved from the Protein Data Bank (PDB). The corresponding sequence matched the UniProt entry Q13148. Protein preparation was carried out using the Protein Preparation Wizard module in Schrödinger Maestro (v14.2), which involved assignment of bond orders, addition of hydrogen atoms, and adjustment of tautomeric and ionization states. Protonation states were assigned to reflect a physiological pH of 7.4, and energy minimization was performed using the OPLS_2005 force field to relieve steric clashes and optimize the geometry. For ligand preparation, 2D structures of Anle138b, ATC141, and ATC142 were converted into 3D conformers using the LigPrep module (Maestro v14.2). The process included generation of multiple tautomeric and ionization states at pH 7.4, removal of salts, and energy minimization using the OPLS_2005 force field with an RMSD convergence threshold of 0.01 Å. Putative ligand-binding sites were identified using the SiteMap module (Maestro v14.2). Molecular docking was performed using the Glide module in standard precision (SP) mode with the OPLS_2005 force field, where the protein was kept rigid and ligands were docked flexibly. Binding affinities of the ligand-protein complexes were further evaluated using Prime MM-GBSA calculations (Maestro v14.2), based on the VSGB solvation model and OPLS_2005 force field. The binding free energy (ΔG_bind) was calculated using the following formula: ΔG_bind = ΔG_solvation + ΔE_minimized + ΔG_surface. Ligand-protein interactions were visualized using the Ligand Interaction Diagram tool to generate 2D interaction maps.

### Statistics analysis

The data are presented as the mean (±SEM) of at least two independent experiments. For data analysis, GraphPad Prism 8 software was used. Statistics were performed using two-tailed unpaired Student’s *t*-test (degree of freedom = n-1). Survival curves were generated using the Kaplan-Meier method and compared using the log-rank (Mantel-Cox) test. Values were considered significant for *p<0.05, **p<0.01, ***p<0.001, and ****p<0.0001.

## List of Supplementary Materials

Fig. S1 to S8 for multiple supplementary figures

Table S1 and S2 for multiple supplementary tables

Methods

## Acknowledgments

We thank the patients and families for donating samples for research and the clinicians and research staff for assistance in sample collection, processing, and curation. We also thank K. C. Park and the members of LNP Solution for the help with in silico analyses.

## Authors’ Contributions

Y.T.K., D.Y.P., and H.Y.K. conceived and designed the experimental approach. D.Y.P., H.Y.K., E.H.C., G.E.L. performed cellular experiments and statistical analyses. D.Y.P., S.H.K., Hye Yeon Kim performed all animal experiments and statistical analyses. J.A.L. and H.N.C. conducted motor neuron differentiation and ALS patient-derived iPSC experiments. Y.H.S. and D.H.P. produced the lentivirus used for primary neuron experiments. Y.T.K., D.Y.P., E.H.C. prepared the manuscript with input from all authors.

## Data and Materials Availability

All data supporting the findings of this study are available within the paper and its supplementary materials. Additional materials and protocols may be requested from the authors.

## References

1. P. Rojas, A. I. Ramírez, J. A. Fernández-Albarral, I. López-Cuenca, E. Salobrar-García, M. Cadena, L. Elvira-Hurtado, J. J. Salazar, R. de Hoz, J. M. Ramírez, Amyotrophic lateral sclerosis: a neurodegenerative motor neuron disease with ocular involvement. Frontiers in Neuroscience 14, 566858 (2020).

2. S. Niedermeyer, M. Murn, P. J. Choi, Respiratory failure in amyotrophic lateral sclerosis. Chest 155, 401–408 (2019).

3. J. Barberio, C. Lally, V. Kupelian, O. Hardiman, W. D. Flanders, Estimated familial amyotrophic lateral sclerosis proportion: a literature review and meta-analysis. Neurology: Genetics 9, e200109 (2023).

4. R. J. Mead, N. Shan, H. J. Reiser, F. Marshall, P. J. Shaw, Amyotrophic lateral sclerosis: a neurodegenerative disorder poised for successful therapeutic translation. Nature Reviews Drug Discovery 22, 185–212 (2023).

5. L. B. Tovar-y-Romo, L. D. Santa-Cruz, A. Zepeda, R. Tapia, Chronic elevation of extracellular glutamate due to transport blockade is innocuous for spinal motoneurons in vivo. Neurochemistry international 54, 186–191 (2009).

6. Y. Yoshii, A. Otomo, L. Pan, M. Ohtsuka, S. Hadano, Loss of glial fibrillary acidic protein marginally accelerates disease progression in a SOD1H46R transgenic mouse model of ALS. Neuroscience research 70, 321–329 (2011).

7. E. N. Guerrero, H. Wang, J. Mitra, P. M. Hegde, S. E. Stowell, N. F. Liachko, B. C. Kraemer, R. M. Garruto, K. Rao, M. L. Hegde, TDP-43/FUS in motor neuron disease: Complexity and challenges. Progress in neurobiology 145, 78–97 (2016).

8. E. Parobkova, R. Matej, Amyotrophic lateral sclerosis and frontotemporal lobar degenerations: similarities in genetic background. Diagnostics 11, 509 (2021).

9. H. P. Nguyen, C. Van Broeckhoven, J. van der Zee, ALS genes in the genomic era and their implications for FTD. Trends in Genetics 34, 404–423 (2018).

10. S. May, D. Hornburg, M. H. Schludi, T. Arzberger, K. Rentzsch, B. M. Schwenk, F. A. Grässer, K. Mori, E. Kremmer, J. Banzhaf-Strathmann, C9orf72 FTLD/ALS-associated Gly-Ala dipeptide repeat proteins cause neuronal toxicity and Unc119 sequestration. Acta neuropathologica 128, 485–503 (2014).

11. N. E. Farrawell, I. A. Lambert-Smith, S. T. Warraich, I. P. Blair, D. N. Saunders, D. M. Hatters, J. J. Yerbury, Distinct partitioning of ALS associated TDP-43, FUS and SOD1 mutants into cellular inclusions. Scientific reports 5, 13416 (2015).

12. I. Casafont, R. Bengoechea, O. Tapia, M. T. Berciano, M. Lafarga, TDP-43 localizes in mRNA transcription and processing sites in mammalian neurons. Journal of structural biology 167, 235–241 (2009).

13. C. F. Sephton, B. Cenik, B. K. Cenik, J. Herz, G. Yu, TDP-43 in central nervous system development and function: clues to TDP-43-associated neurodegeneration. Biological chemistry 393, 589–594 (2012).

14. F. Arnold, A. Nguyen, R. Bedlack, C. Bennett, A. La Spada, Intercellular transmission of pathogenic proteins in ALS: Exploring the pathogenic wave. Neurobiology of Disease, 106218 (2023).

15. B. A. Berning, A. K. Walker, The pathobiology of TDP-43 C-terminal fragments in ALS and FTLD. Frontiers in neuroscience 13, 335 (2019).

16. A. Prasad, V. Bharathi, V. Sivalingam, A. Girdhar, B. K. Patel, Molecular mechanisms of TDP-43 misfolding and pathology in amyotrophic lateral sclerosis. Frontiers in molecular neuroscience 12, 25 (2019).

17. S. Dhakal, C. E. Wyant, H. E. George, S. E. Morgan, V. Rangachari, Prion-like C-terminal domain of TDP-43 and α-Synuclein interact synergistically to generate neurotoxic hybrid fibrils. Journal of molecular biology 433, 166953 (2021).

18. E. Pokrishevsky, L. I. Grad, N. R. Cashman, TDTDP-43 or FUS-induced misfolded human wild-type SOD1 can propagate intercellularly in a prion-like fashionhion. Scientific reports 6, 22155 (2016).

19. T. R. Suk, M. W. Rousseaux, The role of TDP-43 mislocalization in amyotrophic lateral sclerosis. Molecular neurodegeneration 15, 1–16 (2020).

20. M. Jo, S. Lee, Y.-M. Jeon, S. Kim, Y. Kwon, H.-J. Kim, The role of TDP-43 propagation in neurodegenerative diseases: integrating insights from clinical and experimental studies. Experimental & molecular medicine 52, 1652–1662 (2020).

21. K. E. Prater, C. S. Latimer, S. Jayadev, Glial TDP-43 and TDP-43 induced glial pathology, focus on neurodegenerative proteinopathy syndromes. Glia 70, 239–255 (2022).

22. P. Soares, C. Silva, D. Chavarria, F. S. Silva, P. J. Oliveira, F. Borges, Drug discovery and amyotrophic lateral sclerosis: emerging challenges and therapeutic opportunities. Ageing research reviews 83, 101790 (2023).

23. Y. Saitoh, Y. Takahashi, Riluzole for the treatment of amyotrophic lateral sclerosis. Neurodegenerative disease management 10, 343–355 (2020).

24. H. Cho, S. Shukla, Role of edaravone as a treatment option for patients with amyotrophic lateral sclerosis. Pharmaceuticals 14, 29 (2020).

25. H. Moriyama, T. Yokota, Recent Progress of Antisense Oligonucleotide Therapy for Superoxide-Dismutase-1-Mutated Amyotrophic Lateral Sclerosis: Focus on Tofersen. Genes 15, 1342 (2024).

26. J. A. Gregory, C. M. Hickey, J. Chavez, A. M. Cacace, New therapies on the horizon: Targeted protein degradation in neuroscience. Cell Chemical Biology 31, 1688–1698 (2024).

27. M. Békés, D. R. Langley, C. M. Crews, PROTAC targeted protein degraders: the past is prologue. Nature Reviews Drug Discovery 21, 181–200 (2022).

28. Y.-L. Tseng, P.-C. Lu, C.-C. Lee, R.-Y. He, Y.-A. Huang, Y.-C. Tseng, T.-J. R. Cheng, J. J.-T. Huang, J.-M. Fang, Degradation of neurodegenerative disease-associated TDP-43 aggregates and oligomers via a proteolysis-targeting chimera. Journal of biomedical science 30, 27 (2023).

29. D. M. Rubin, D. Finley, Proteolysis: The proteasome: a protein-degrading organelle? Current Biology 5, 854–858 (1995).

30. T.-M. Sonninen, G. Goldsteins, N. Laham-Karam, J. Koistinaho, Š. Lehtonen, Proteostasis disturbances and inflammation in neurodegenerative diseases. Cells 9, 2183 (2020).

31. A. Ciechanover, Y. T. Kwon, Degradation of misfolded proteins in neurodegenerative diseases: therapeutic targets and strategies. Experimental & molecular medicine 47, e147–e147 (2015).

32. A. Ciechanover, Y. T. Kwon, Protein quality control by molecular chaperones in neurodegeneration. Frontiers in neuroscience 11, 185 (2017).

33. L. Zhao, J. Zhao, K. Zhong, A. Tong, D. Jia, Targeted protein degradation: mechanisms, strategies and application. Signal Transduction and Targeted Therapy 7, 113 (2022).

34. N. B. Nedelsky, P. K. Todd, J. P. Taylor, Autophagy and the ubiquitin-proteasome system: collaborators in neuroprotection. Biochimica et Biophysica Acta (BBA)-Molecular Basis of Disease 1782, 691–699 (2008).

35. E. L. Scotter, C. Vance, A. L. Nishimura, Y.-B. Lee, H.-J. Chen, H. Urwin, V. Sardone, J. C. Mitchell, B. Rogelj, D. C. Rubinsztein, Differential roles of the ubiquitin proteasome system and autophagy in the clearance of soluble and aggregated TDP-43 species. Journal of cell science 127, 1263–1278 (2014).

36. M. Budini, E. Buratti, E. Morselli, A. Criollo, Autophagy and its impact on neurodegenerative diseases: new roles for TDP-43 and C9orf72. Frontiers in Molecular Neuroscience 10, 170 (2017).

37. A. Fleming, M. Bourdenx, M. Fujimaki, C. Karabiyik, G. J. Krause, A. Lopez, A. Martín-Segura, C. Puri, A. Scrivo, J. Skidmore, The different autophagy degradation pathways and neurodegeneration. Neuron 110, 935–966 (2022).

38. A. Varshavsky, N-degron pathways. Proceedings of the National Academy of Sciences 121, e2408697121 (2024).

39. A. J. Heo, S. B. Kim, Y. T. Kwon, C. H. Ji, The N-degron pathway: From basic science to therapeutic applications. Biochimica et Biophysica Acta (BBA)-Gene Regulatory Mechanisms, 194934 (2023).

40. Y. T. Kwon, Z. Xia, I. V. Davydov, S. H. Lecker, A. Varshavsky, Construction and analysis of mouse strains lacking the ubiquitin ligase UBR1 (E3α) of the N-end rule pathway. Molecular and cellular biology 21, 8007–8021 (2001).

41. T. Tasaki, L. C. Mulder, A. Iwamatsu, M. J. Lee, I. V. Davydov, A. Varshavsky, M. Muesing, Y. T. Kwon, A family of mammalian E3 ubiquitin ligases that contain the UBR box motif and recognize N-degrons. Molecular and cellular biology 25, 7120–7136 (2005).

42. S. M. Sriram, R. Banerjee, R. S. Kane, Y. T. Kwon, Multivalency-assisted control of intracellular signaling pathways: application for ubiquitin-dependent N-end rule pathway. Chemistry & biology 16, 121–131 (2009).

43. H. Cha-Molstad, J. E. Yu, Z. Feng, S. H. Lee, J. G. Kim, P. Yang, B. Han, K. W. Sung, Y. D. Yoo, J. Hwang, p62/SQSTM1/Sequestosome-1 is an N-recognin of the N-end rule pathway which modulates autophagosome biogenesis. Nature communications 8, 102 (2017).

44. H. Cha-Molstad, K. S. Sung, J. Hwang, K. A. Kim, J. E. Yu, Y. D. Yoo, J. M. Jang, D. H. Han, M. Molstad, J. G. Kim, Amino-terminal arginylation targets endoplasmic reticulum chaperone BiP for autophagy through p62 binding. Nature cell biology 17, 917–929 (2015).

45. Y. D. Yoo, S. R. Mun, C. H. Ji, K. W. Sung, K. Y. Kang, A. J. Heo, S. H. Lee, J. Y. An, J. Hwang, X.-Q. Xie, N-terminal arginylation generates a bimodal degron that modulates autophagic proteolysis. Proceedings of the National Academy of Sciences 115, E2716–E2724 (2018).

46. C. H. Ji, H. Y. Kim, M. J. Lee, A. J. Heo, D. Y. Park, S. Lim, S. Shin, S. Ganipisetti, W. S. Yang, C. A. Jung, The AUTOTAC chemical biology platform for targeted protein degradation via the autophagy-lysosome system. Nature communications 13, 904 (2022).

47. H. Cha-Molstad, J. E. Yu, S. H. Lee, J. G. Kim, K. S. Sung, J. Hwang, Y. D. Yoo, Y. J. Lee, S. T. Kim, D. H. Lee, Modulation of SQSTM1/p62 activity by N-terminal arginylation of the endoplasmic reticulum chaperone HSPA5/GRP78/BiP. Autophagy 12, 426–428 (2016).

48. C. H. Ji, H. Y. Kim, A. J. Heo, S. H. Lee, M. J. Lee, S. B. Kim, G. Srinivasrao, S. R. Mun, H. Cha-Molstad, A. Ciechanover, The N-degron pathway mediates ER-phagy. Molecular cell 75, 1058–1072. e1059 (2019).

49. S. M. Shim, H. R. Choi, S. C. Kwon, H. Y. Kim, K. W. Sung, E. J. Jung, S. R. Mun, T. H. Bae, D. H. Kim, Y. S. Son, The Cys-N-degron pathway modulates pexophagy through the N-terminal oxidation and arginylation of ACAD10. Autophagy 19, 1642–1661 (2023).

50. E. J. Jung, K. W. Sung, T. H. Bae, H.-Y. Kim, H. R. Choi, S. H. Kim, C. H. Jung, S. R. Mun, Y. S. Son, S. Kim, The N-degron pathway mediates lipophagy: the chemical modulation of lipophagy in obesity and NAFLD. Metabolism 146, 155644 (2023).

51. J. Lee, K. W. Sung, E.-J. Bae, D. Yoon, D. Kim, J. S. Lee, D.-h. Park, D. Y. Park, S. R. Mun, S. C. Kwon, Targeted degradation of⍺-synuclein aggregates in Parkinson’s disease using the AUTOTAC technology. Molecular Neurodegeneration 18, 1–21 (2023).

52. T. H. Bae, K. W. Sung, T. M. Pham, A. J. Najy, A. Zamiri, H. Jang, S. R. Mun, S. Kim, H. K. Kwon, Y. S. Son, An Autophagy-Targeting Chimera Induces Degradation of Androgen Receptor Mutants and AR-v7 in Castration-Resistant Prostate Cancer. Cancer Research 85, 342–359 (2025).

53. J. Shenoy, A. Lends, M. Berbon, M. Bilal, N. El Mammeri, M. Bertoni, A. Saad, E. Morvan, A. Grélard, S. Lecomte, Structural polymorphism of the low-complexity C-terminal domain of TDP-43 amyloid aggregates revealed by solid-state NMR. Frontiers in Molecular Biosciences 10, 1148302 (2023).

54. L.-L. Jiang, M.-X. Che, J. Zhao, C.-J. Zhou, M.-Y. Xie, H.-Y. Li, J.-H. He, H.-Y. Hu, Structural transformation of the amyloidogenic core region of TDP-43 protein initiates its aggregation and cytoplasmic inclusion. Journal of Biological Chemistry 288, 19614–19624 (2013).

55. M. Y. Fang, S. Markmiller, A. Q. Vu, A. Javaherian, W. E. Dowdle, P. Jolivet, P. J. Bushway, N. A. Castello, A. Baral, M. Y. Chan, Small-molecule modulation of TDP-43 recruitment to stress granules prevents persistent TDP-43 accumulation in ALS/FTD. Neuron 103, 802–819. e811 (2019).

56. R. L. French, A. N. Reeb, H. Aligireddy, N. Kedia, D. D. Dhavale, Z. R. Grese, P. T. Kotzbauer, J. Bieschke, Y. M. Ayala, TDP-43 oligomers detected as initial intermediate species during aggregate formation. bioRxiv, 499343 (2018).

57. R. H. Baloh, TDP-43: the relationship between protein aggregation and neurodegeneration in amyotrophic lateral sclerosis and frontotemporal lobar degeneration. The FEBS journal 278, 3539–3549 (2011).

58. Z.-W. Du, H. Chen, H. Liu, J. Lu, K. Qian, C.-L. Huang, X. Zhong, F. Fan, S.-C. Zhang, Generation and expansion of highly pure motor neuron progenitors from human pluripotent stem cells. Nature communications 6, 6626 (2015).

59. J. Gao, L. Wang, M. L. Huntley, G. Perry, X. Wang, Pathomechanisms of TDP-43 in neurodegeneration. Journal of neurochemistry 146, 7–20 (2018).

60. Y. T. Kwon, C. H. Ji, S. Ganipisetti, H. Y. Kim, S. R. Mun, C. H. Jung, E. J. Jung, K. W. Sung, Novel AUTOTAC Chimeric Compounds and Compositions for the Prevention, Amelioration, or Treatment of Diseases via Targeted Protein Degradation. WO patent 2020/022785 A1, (2020).

61. I. Wegorzewska, S. Bell, N. J. Cairns, T. M. Miller, R. H. Baloh, TDP-43 mutant transgenic mice develop features of ALS and frontotemporal lobar degeneration. Proceedings of the National Academy of Sciences 106, 18809–18814 (2009).

62. L. Bargsted, D. B. Medinas, F. Martínez Traub, P. Rozas, N. Muñoz, M. Nassif, C. Jerez, A. Catenaccio, F. A. Court, C. Hetz, Disulfide cross-linked multimers of TDP-43 and spinal motoneuron loss in a TDP-43A315T ALS/FTD mouse model. Scientific Reports 7, 14266 (2017).

63. D. D. Biswas, R. Sethi, Y. Woldeyohannes, E. R. Scarrow, L. El Haddad, J. Lee, M. K. ElMallah, Respiratory pathology in the TDP-43 transgenic mouse model of amyotrophic lateral sclerosis. Frontiers in Physiology 15, 1430875 (2024).

64. M. Tsekrekou, M. Giannakou, K. Papanikolopoulou, G. Skretas, Protein aggregation and therapeutic strategies in SOD1-and TDP-43-linked ALS. Frontiers in Molecular Biosciences 11, 1383453 (2024).

65. M. K. Jaiswal, Riluzole and edaravone: A tale of two amyotrophic lateral sclerosis drugs. Medicinal Research Reviews 39, 733–748 (2019).

66. X. Zhang, L. Li, S. Chen, D. Yang, Y. Wang, X. Zhang, Z. Wang, W. Le, Rapamycin treatment augments motor neuron degeneration in SOD1G93A mouse model of amyotrophic lateral sclerosis. Autophagy 7, 412–425 (2011).

67. I.-F. Wang, B.-S. Guo, Y.-C. Liu, C.-C. Wu, C.-H. Yang, K.-J. Tsai, C.-K. J. Shen, Autophagy activators rescue and alleviate pathogenesis of a mouse model with proteinopathies of the TAR DNA-binding protein 43. Proceedings of the National Academy of Sciences 109, 15024–15029 (2012).

68. A. Bhattacharya, A. Bokov, F. L. Muller, A. L. Jernigan, K. Maslin, V. Diaz, A. Richardson, H. Van Remmen, Dietary restriction but not rapamycin extends disease onset and survival of the H46R/H48Q mouse model of ALS. Neurobiology of aging 33, 1829–1832 (2012).

69. X. G. Chen, F. Liu, X. F. Song, Z. H. Wang, Z. Q. Dong, Z. Q. Hu, R. Z. Lan, W. Guan, T. G. Zhou, X. M. Xu, Rapamycin regulates Akt and ERK phosphorylation through mTORC1 and mTORC2 signaling pathways. Molecular Carcinogenesis: Published in cooperation with the University of Texas MD Anderson Cancer Center 49, 603–610 (2010).

70. C. Gomes, C. Escrevente, J. Costa, Mutant superoxide dismutase 1 overexpression in NSC-34 cells: effect of trehalose on aggregation, TDP-43 localization and levels of co-expressed glycoproteins. Neuroscience letters 475, 145–149 (2010).

71. Y. Li, Y. Guo, X. Wang, X. Yu, W. Duan, K. Hong, J. Wang, H. Han, C. Li, Trehalose decreases mutant SOD1 expression and alleviates motor deficiency in early but not end-stage amyotrophic lateral sclerosis in a SOD1-G93A mouse model. Neuroscience 298, 12–25 (2015).

72. N. D. Perera, R. K. Sheean, C. L. Lau, Y. S. Shin, P. M. Beart, M. K. Horne, B. J. Turner, Rilmenidine promotes MTOR-independent autophagy in the mutant SOD1 mouse model of amyotrophic lateral sclerosis without slowing disease progression. Autophagy 14, 534–551 (2018).

73. N. D. Perera, D. Tomas, N. Wanniarachchillage, B. Cuic, S. J. Luikinga, V. Rytova, B. J. Turner, Stimulation of mTOR-independent autophagy and mitophagy by rilmenidine exacerbates the phenotype of transgenic TDP-43 mice. Neurobiology of Disease 154, 105359 (2021).

74. H. Beck, M. Härter, B. Haß, C. Schmeck, L. Baerfacker, Small molecules and their impact in drug discovery: A perspective on the occasion of the 125th anniversary of the Bayer Chemical Research Laboratory. Drug Discovery Today 27, 1560–1574 (2022).

