## Supplementary Materials for "Targeted degradation of pathogenic TDP-43 proteins in amyotrophic lateral sclerosis using the AUTOTAC platform"

**The PDF file includes:**

Fig. S1. Structure and synthesis of AUTOTACs
Fig. S2. Spectroscopic and analytical characterization of ATC142
Fig. S3. Screening of degradative efficacy of chemical ligands and AUTOTACs
Fig. S4. TDP-43 overexpression induces ALS-related phenotypes
Fig. S5. Generation and characterization of ALS patient-derived iPSC using fibroblast cells
Fig. S6. Differentiation of human iPSCs into cholinergic motor neurons
Fig. S7. ATC141 selectively degrades pathological TDP-43 in vitro
Fig. S8. Comparative analysis of motor function and survival in non-transgenic and TDP-43 A315T mice
Table S1. In silico analysis of predicted binding sites and docking scores for Anle138b
Table S2. Pharmacokinetic (PK) parameters of ATC142
Supplementary Methods: Chemical synthesis of ATC142

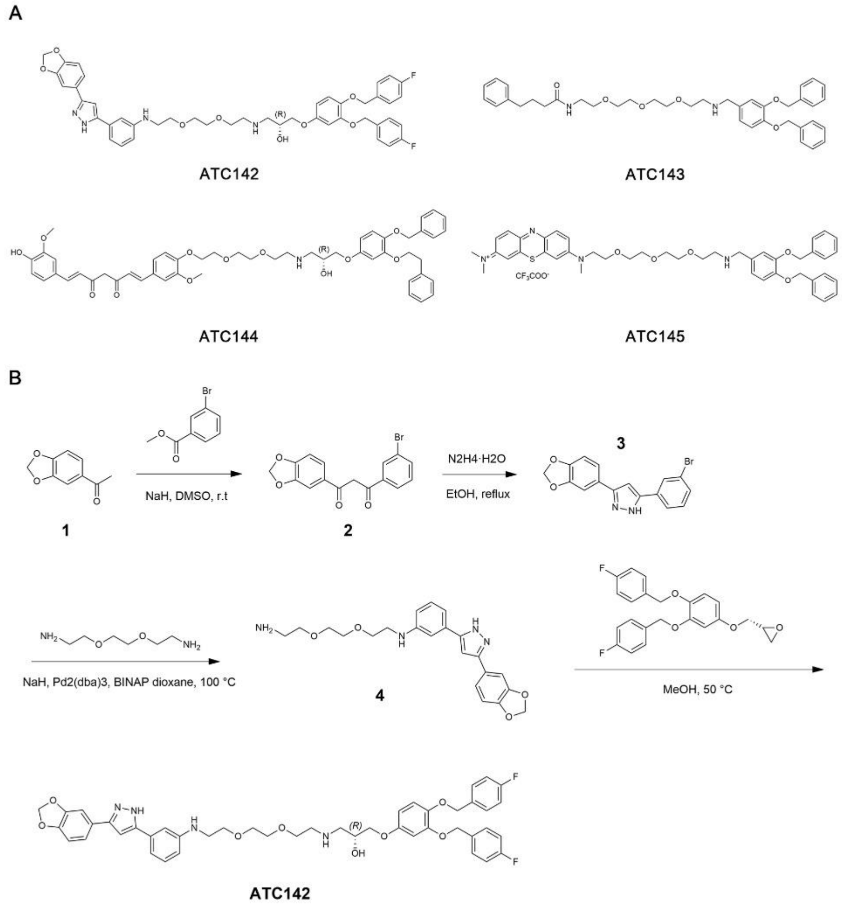


**Supplementary Fig. 1. Structure and synthesis of AUTOTACs**

**(A)** Chemical structures of AUTOTAC compounds containing TBL moieties, comprising Anle138b (ATC142), 4-PBA (ATC143), curcumin (ATC144), or methylene blue (ATC145). (**B)** Synthesis scheme for ATC142.

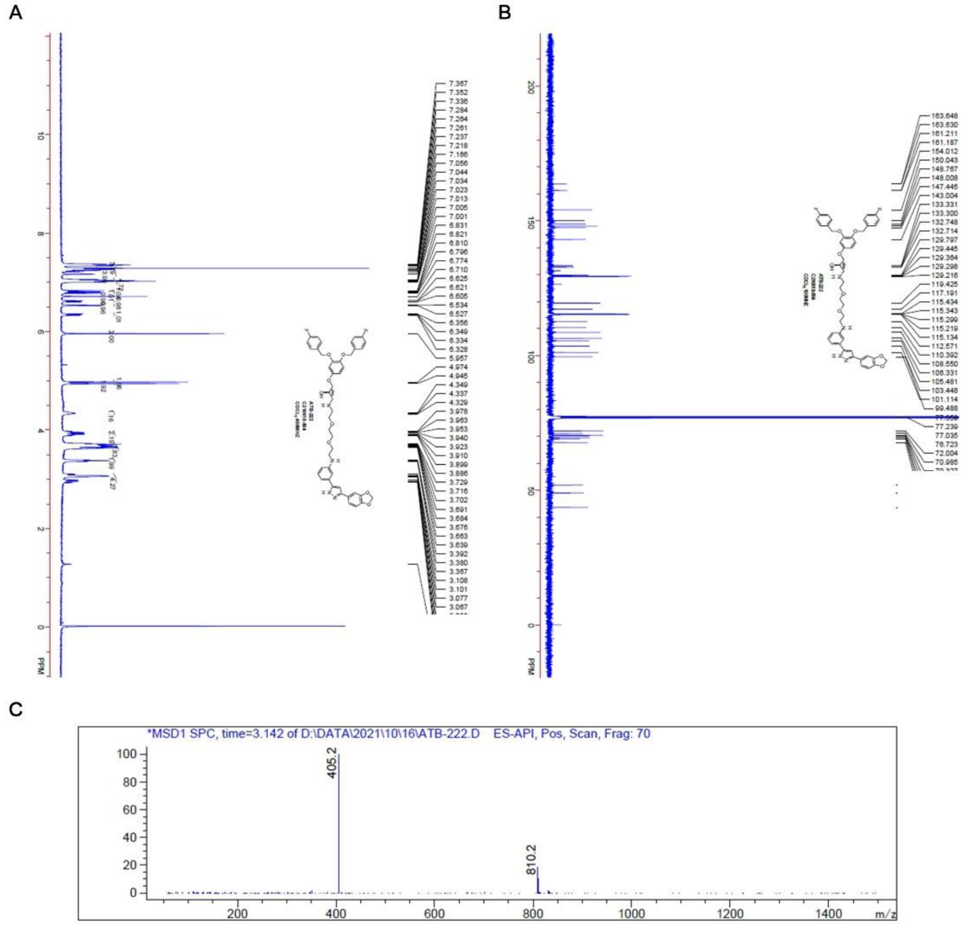


**Supplementary Fig. 2. Spectroscopic and analytical characterization of ATC142**

**(A)** $^{1}$H-NMR spectrum of ATC142. **(B)** $^{13}$C-NMR spectrum of ATC142. (**C)** LC-MS chromatogram of ATC142.

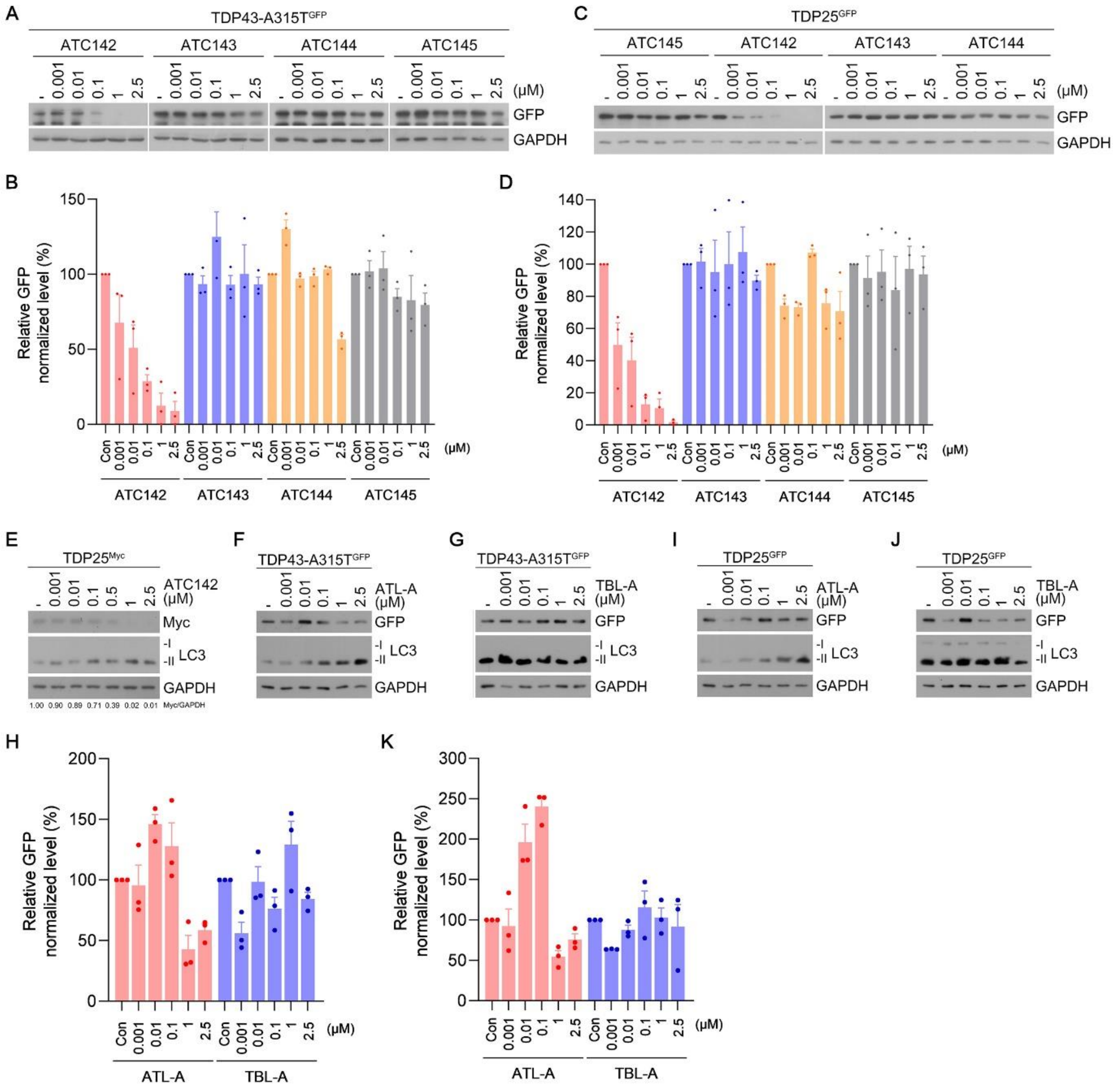


**Supplementary Fig. 3. Screening of degradative efficacy of chemical ligands and AUTOTACs**

**(A)** Immunoblotting analysis of HEK293T cells transfected with TDP-43 A315T-GFP and treated with ATC142, ATC143, ATC144, or ATC145 at the indicated concentrations (24 h). (**B)** Quantification of GFP levels shown in (A). **(C)** Identical to (A), but in HEK293T cells transfected with TDP25-GFP. **(D)** Quantification of GFP levels shown in (C). **(E)** Immunoblotting analysis of HeLa cells transfected with TDP25-Myc and treated with ATC142 at the indicated concentrations (24 h). **(F)** Immunoblotting analysis of HeLa cells transfected with TDP-43 A315T-GFP and treated with ATL-A at the indicated concentrations (24 h). **(G)** Identical to (F), but treated with TBL-A (Anle138b) at the indicated concentrations (24 h). **(H)** Quantification of GFP levels shown in (F) and (G). **(I)** Immunoblotting analysis of HeLa cells transfected with TDP25-GFP and treated with ATL-A at the indicated concentrations (24 h). **(J)** Identical to (I), but treated with TBL-A (Anle138b) at the indicated concentrations (24 h). **(K)** Quantification of GFP levels shown in (I) and (J).

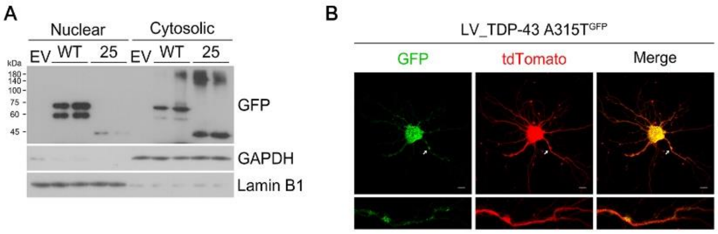


**Supplementary Fig. 4. TDP-43 overexpression induces ALS-related phenotypes**

**(A)** Nuclear/cytosolic fractionation assay in HEK293T cells transfected with either TDP-43 WT-GFP or TDP25-GFP. **(B)** Immunocytochemistry of primary hippocampal neurons transduced with TDP-43 A315T-GFP lentivirus. Scale bar, 10 µm.

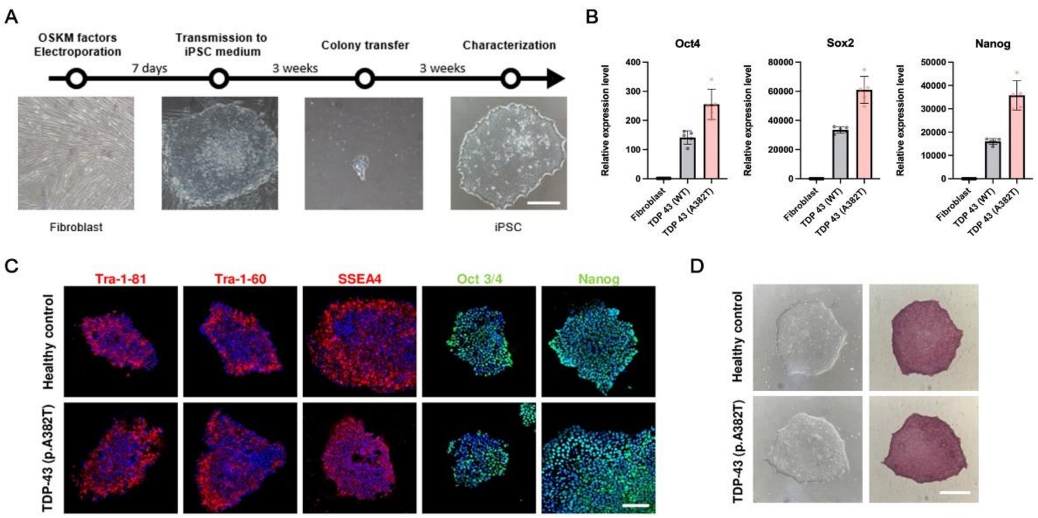


**Supplementary Fig. 5. Generation and characterization of ALS patient-derived iPSC using fibroblast cells**

**(A)** Schematic representation of the iPSC generation workflow. Scale bar, 100 µm. **(B)** Real-time PCR analysis of pluripotency marker expression (Oct3/4, Sox2 and Nanog) in ALS iPSCs. **(C)** Immunostaining of iPSC colonies with pluripotency markers. Scale bar, 100 µm. **(D)** Morphology and alkaline phosphatase staining of iPSC colonies. Scale bar, 100 µm.

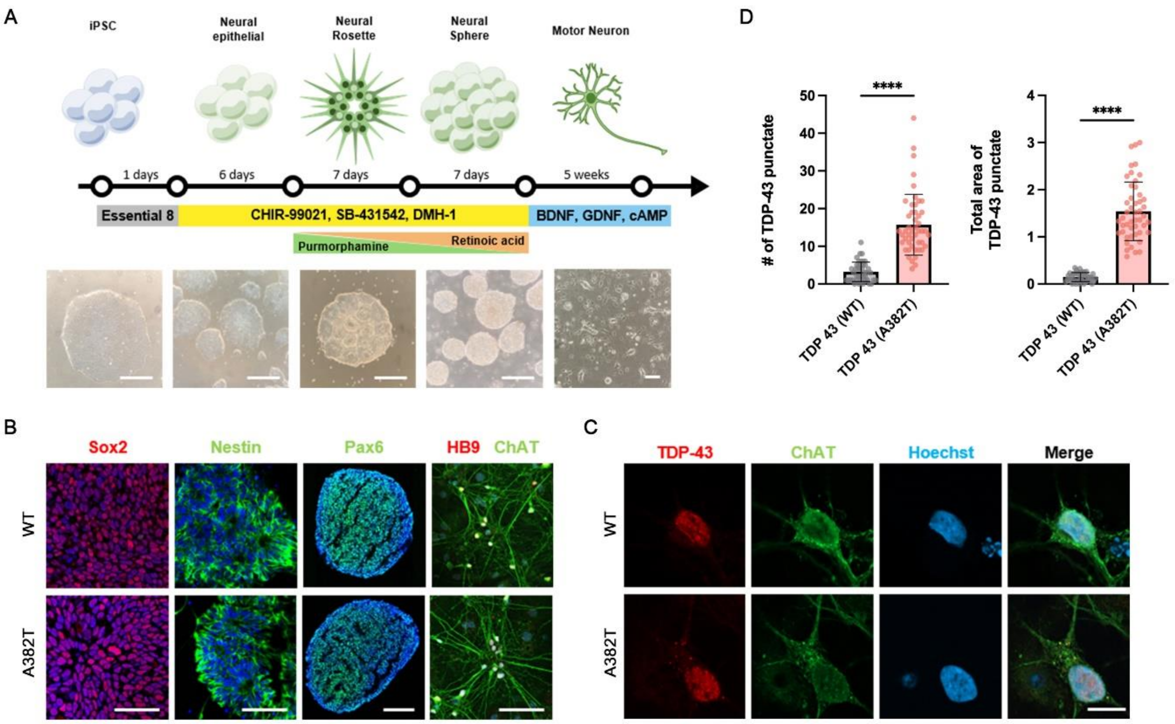


**Supplementary Fig. 6. Differentiation of human iPSCs into cholinergic motor neurons**

**(A)** Schematic representation of the cholinergic motor neuron differentiation protocol, including media supplements, growth factors, and small molecules. **(B)** Representative immunocytochemical images showing motor neural differentiation. (SOX2, Nestin, HB9: scale bars: 50 µm; Pax6: scale bar, 200 µm) **(C)** Cytosolic TDP-43 aggregates observed in motor neurons derived from ALS patient iPSCs. Scale bar: 10 µm **(D)** Quantification of the number and size of cytosolic TDP-43 aggregates shown in (C). Data are presented as mean ± S.E.M. Statistical significance is indicated as follows: ****P ≤ 0.0001. (Student's *t*-test).

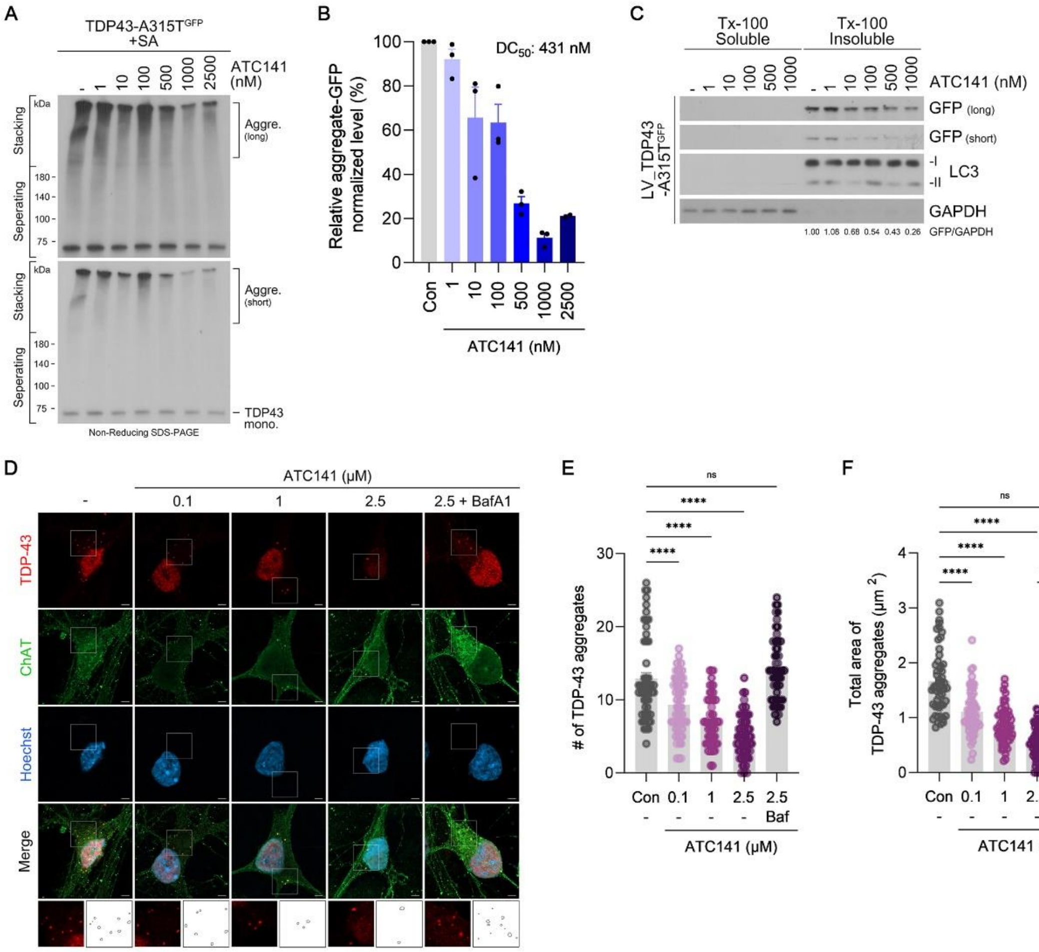

**Supplementary Fig. 7. ATC141 selectively degrades pathological TDP-43 in vitro**

**(A)** Non-reducing SDS PAGE of HEK293T cells transfected with TDP-43 A315T-GFP and treated with sodium arsenite (200 µM, 24 h). ATC141 was applied at the indicated concentrations (24 h), and GFP-tagged proteins were detected. **(B)** Quantification of aggregate-sized GFP retained in the stacking gel shown in (A). **(C)** Triton X-100 fractionation assay of primary cortical neurons transduced with TDP-43 A315T-GFP lentivirus and treated with ATC141 at the indicated concentrations (24 h). **(D)** Immunocytochemistry of ALS iPSCs treated with ATC141 at the indicated concentrations (24 h), with or without bafilomycin A1 (10 nM, 24 h). **(E)** Quantification of the number of TDP-43 aggregates in (D). **(F)** Quantification of the total area of TDP-43 aggregates in (D). Data are presented as mean ± S.E.M. Statistical significance is indicated as follows: ns, not significant; ****P ≤ 0.0001. (Student's *t*-test).

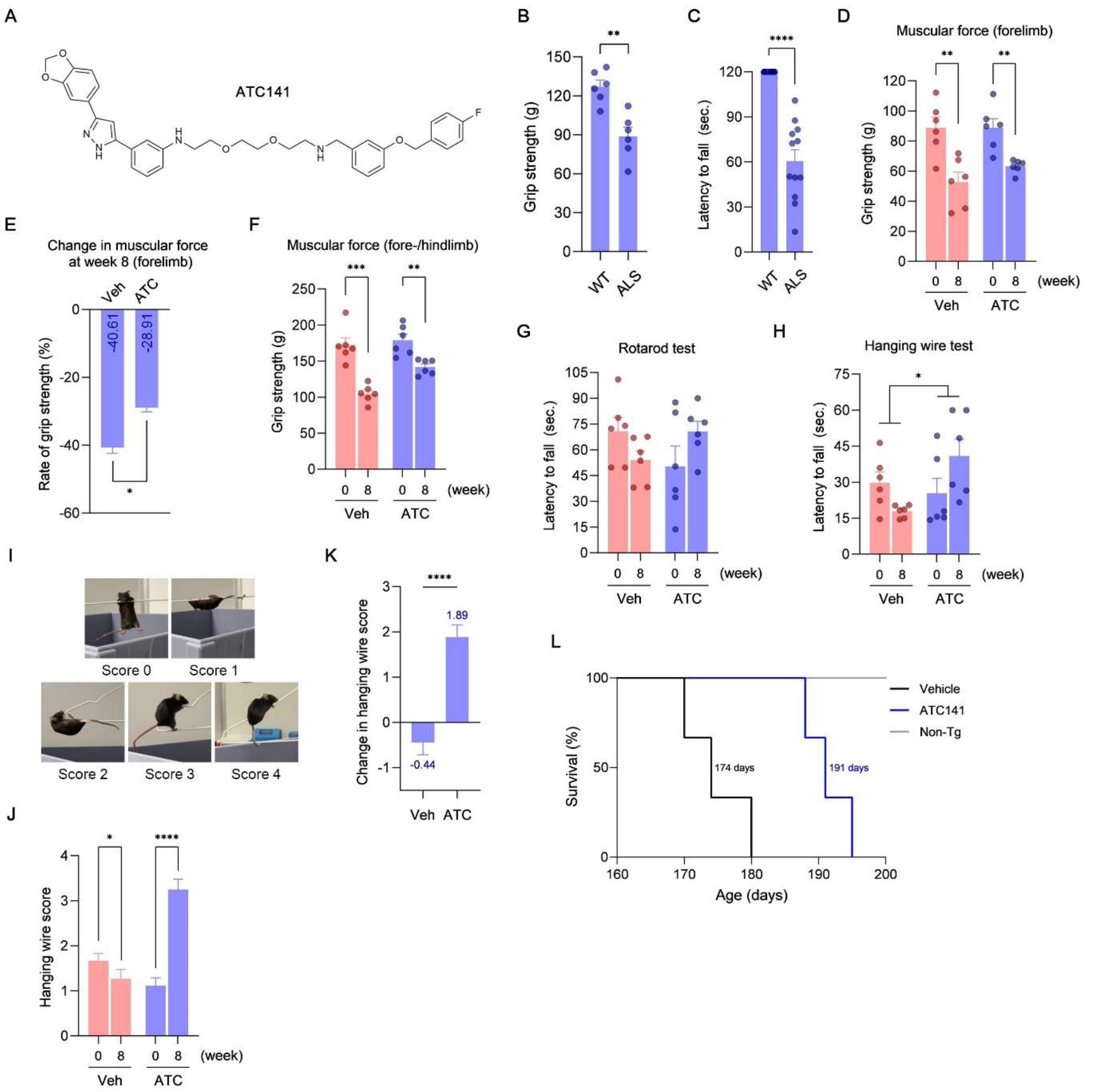


**Supplementary Fig. 8. Comparative analysis of motor function and survival in non-transgenic and TDP-43 A315T mice**

**(A)** Chemical structure of ATC141. **(B)** Comparative analysis of forelimb grip strength at week 0 between non-transgenic wild-type and TDP-43 A315T mice. **(C)** Comparative analysis of latency to fall in the rotarod test at week 0 between non-transgenic wild-type and TDP-43 A315T mice. **(D)** Forelimb grip strength of TDP-43 A315T mice treated with vehicle or ATC141, measured at week 0 and week 8. **(E)** Comparative analysis of forelimb grip strength at week 8 between vehicle- and ATC141-treated TDP-43 A315T mice. **(F)** Combined forelimb and hindlimb grip strength of TDP-43 A315T mice treated with vehicle or ATC141, measured at weeks 0 and 8. **(G)** Latency to fall in the rotarod test of TDP-43 A315T mice treated with vehicle or ATC141, measured at weeks 0 and 8. **(H)** Latency to fall in the hanging wire test of TDP-43 A315T mice treated with vehicle or ATC141, measured at weeks 0 and 8. **(I)** Representative images illustrating success assessment scores in the hanging wire test. **(J)** Average success assessment scores from the hanging wire test in vehicle- and ATC141-injected mice, measured at weeks 0 and 8. **(K)** Comparative analysis of success assessment scores at week 8 between vehicle- and ATC141-injected TDP-43 A315T mice in the hanging wire test. **(L)** Survival rates (%) of non-transgenic, and vehicle-treated and ATC141-treated TDP-43 A315T mice over the study period. Data are presented as mean ± S.E.M. Survival was analyzed by Kaplan-Meier method and log-rank test. Statistical significance is indicated as follows: *P ≤ 0.05; **P ≤ 0.01; ***P ≤ 0.001; ****P ≤ 0.0001. (Student's *t*-test).

| Site No. | Site Volume (Å³) | Site Score | Dscore | Key Residues | Glide Score (kcal/mol) | MM-GBSA ΔG (kcal/mol) |
|---|---|---|---|---|---|---|
| 1 | 81.97 | 0.59 | 0.52 | N302, A326, S333 | -4.32 | -44.45 |
| 2 | 108.73 | 0.76 | 0.76 | G335, M337, N343 | -4.29 | -65.40 |
| 3 | 40.47 | 0.89 | 1.03 | N344, G346 | -5.19 | -44.42 |
| 4 | 18.18 | 0.47 | 0.48 | A315, F316, N360 | -3.14 | -24.52 |

**Supplementary Table 1. In silico analysis of predicted binding sites and docking scores for Anle138b**

Summary of computational predictions of ligand-binding sites and docking metrics for Anle138b. The table includes the number, volume, and score of each predicted site, druggability score (Dscore), key interacting residues, glide docking score, and MM-GBSA binding free energy.

PK parameters of ATC142

| Route | IP | SC | PO | IV |
|---|---|---|---|---|
| Dose / Unit (mpk) | 10 | 10 | 10 | 1 |
| $T_{1/2}$ (hr) | 6.2 | 6.1 | 2.3 | 4.7 |
| $T_{max}$ (hr) | 1.4 | 2.3 | 0.8 | 0.1 |
| $C_{max}$ (ng/ml) | 1170.6 | 644.0 | 19.3 | 1284.5 |
| $AUC_{last}$ (ng*h/ml) | 7061.0 | 4771.3 | 48.9 | 499.2 |
| F (%) | | 95.6 | 0.98 | |

**Supplementary Table 2. Pharmacokinetic (PK) parameters of ATC142.**

ATC142 was administered to ICR mice via intraperitoneal (IP), subcutaneous (SC), oral (PO), and intravenous (IV) routes. Pharmacokinetic (PK) parameters, including dose (mg/kg), half-life ($T_{1/2}$), time to maximum concentration ($T_{max}$), maximum plasma concentration ($C_{max}$), area under the plasma concentration-time curve to the last measurable point ($AUC_{last}$), and bioavailability (F, %) were determined.

### SUPPLEMENTARY METHODS

**Chemical synthesis of ATC142**

$^1$H NMR (Fig. S2A) and $^{13}$C NMR (Fig. S2B) spectra were recorded on Bruker Avance III 400 MHz and Bruker Fourier 300 MHz and TMS was used as an internal standard.

LCMS (Fig. S2C) was taken on a quadrupole Mass Spectrometer on Agilent 1260HPLC and 6120MSD (Column: C18 (50 × 4.6 mm, 5 μm) operating in ES (+) or (-) ionization mode; T = 30 $^o$C; flow rate = 1.5 mL/min; detected wavelength: 220 nm, 254nm.

**Scheme 1.** (R)-1-((2-(2-(2-((3-(3-(benzo[d][1,3]dioxol-5-yl)-1H-pyrazol-5-yl)phenyl)amino)ethoxy)ethoxy)ethyl)amino)-3-(3,4-bis((4-fluorobenzyl)oxy)phenoxy)propan-2-ol (**ATC142**; Supplementary Fig. S1A).

1.1 Synthesis of 1-(benzo[d][1,3]dioxol-5-yl)-3-(3-bromophenyl)propane-1,3-dione (**2**)

To a solution of **compound 1** (1-(benzo[d][1,3]dioxol-5-yl)ethan-1-one) (500 g, 3.05 mol, 1.0 eq) and NaH (152 g, 3.81 mol, 1.25 eq, 60% in mineral oil) in dry dimethyl sulfoxide (3000 mL) was stirred at 15 $^o$C for 30 min. Then a solution of methyl 3-bromobenzoate (819 g, 3.81 mol, 1.25 eq) in dimethyl sulfoxide (1500 mL) was added at 20 $^o$C. The resulting mixture was stirred at 25 $^o$C for 2 hrs. The reaction was quenched with saturated aqueous $NH_4Cl$ solution. The mixture was poured into water (10000 mL) and petroleum ether (5000 mL) and then stirred for 30 min at room temperature. Then filtered to give **compound 2 (**1-(benzo[d][1,3]dioxol-5-yl)-3-(3-bromophenyl)propane-1,3-dione) (1000 g, crude) as yellow solid. (TLC: PE/EA=10/1, $R_f$=0.4)

$^1$HNMR ($CDCl_3$, 400 MHz): δ 8.105 (s, 1H), 7.893-7.913 (m, 1H), 7.673-7.693 (m, 1H), 7.618-7.643 (m, 1H), 7.491-7.495 (d, *J* = 1.6 Hz, 1H), 7.362-7.401 (m, 1H), 6.918-6.939 (d, *J* = 8 Hz, 1H), 6.721 (s, 1H), 6.098 (s, 2H)

1.2 Synthesis of 3-(benzo[d][1,3]dioxol-5-yl)-5-(3-bromophenyl)-1H-pyrazole (**3**)

To a solution of **compound 2** (1-(benzo[d][1,3]dioxol-5-yl)-3-(3-bromophenyl)propane-1,3-dione) (1000 g, crude, 2.88 mol, 1.0 eq) and 80% $N_2H_4.H_2O$ (225 g, 3.60 mol, 1.25 eq) in ethanol (15000 mL) was refluxed for 2 hrs. The mixture was cooled to room temperature and filtered to give the **compound 3** (3-(benzo[d][1,3]dioxol-5-yl)-5-(3-bromophenyl)-1H-pyrazole) (600 g, 60.7% from compound 1) as off-white solid. (TLC: PE/EA=5/1, $R_f$=0.3)

$^1$HNMR (DMSO_$d_6$, 400 MHz): δ 13.324 (s, 1H), 8.02 (s, 1H), 7.84 (s, 1H), 7.33-7.52 (m, 4H), 7.21 (s, 1H), 7.03-7.05 (s, 1H), 6.08 (s, 2H)

1.3 Synthesis of N-(2-(2-(2-aminoethoxy)ethoxy)ethyl)-3-(3-(benzo[d][1,3]dioxol-5-yl)-1H-pyrazol-5-yl)aniline (**4**)

To a solution of **compound 3** (3-(benzo[d][1,3]dioxol-5-yl)-5-(3-bromophenyl)-1H-pyrazole) (600 g, 1.75 mol, 1.00 eq) and 2,2'-(ethane-1,2-diylbis(oxy))diethanamine (778 g, 5.25 mol, 3.00 eq) in 1,4-dioxane (6000 mL) was add NaH (210 g, 5.25 mol, 3.00 eq, 60% in mineral oil) slowly at 25 $^o$C. Then added $Pd_2(dba)_3$ (60 g, 0.065 mol, 0.037 eq) and BINAP (120 g, 0.192 mol, 0.11 eq) at 25 $^o$C under $N_2$. The mixture was stirred overnight at 100 $^o$C under $N_2$. The reaction was cooled to room temperature, then quenched with water (12000 mL), extracted with

EA (6000 mL), and concentrated. The crude was purified by column chromatography (DCM/MeOH=50/1~5/1) to give the **compound 4** (N-(2-(2-(2-aminoethoxy)ethoxy)ethyl)-3-(3-(benzo[d][1,3]dioxol-5-yl)-1H-pyrazol-5-yl)aniline) (245 g, 34.1%). (TLC: DCM/MeOH=7/1, $R_f$=0.4)

$^1$HNMR ($CDCl_3$, 400 MHz): δ 7.29-7.25(m, 2H), 7.21-7.23 (m, 1H), 7.07 (s, 1H), 7.00-7.02(m, 1H), 6.83 (d, *J* = 8 Hz, 1H), 6.69 (s, 1H), 6.58 (d, *J* = 8 Hz, 1H), 5.97 (s, 2H), 3.72-3.75 (m, 2H), 3.54-3.67 (m, 6H), 3.32-3.35 (m, 2H), 2.97-2.99 (m, 2H)

1.4 Synthesis of (R)-1-((2-(2-(2-((3-(3-(benzo[d][1,3]dioxol-5-yl)-1H-pyrazol-5-yl)phenyl)amino)ethoxy)ethoxy)ethyl)amino)-3-(3,4-bis((4-fluorobenzyl)oxy)phenoxy)propan-2-ol (**ATC142**)

To a solution of **compound 4** (N-(2-(2-(2-aminoethoxy)ethoxy)ethyl)-3-(3-(benzo[d][1,3]dioxol-5-yl)-1H-pyrazol-5-yl)aniline) (245 g, 597 mmol, 1.00 eq) and (R)-2-((3,4-bis((4-fluorobenzyl)oxy)phenoxy)methyl)oxirane (238 g, 597 mmol, 1.00 eq) in MeOH (4000 mL) was stirred overnight at 50 $^{o}$C. The reaction was concentrated and purified by column chromatography (DCM/MeOH=50/1~20/1) to give the **ATC142** ((R)-1-((2-(2-(2-((3-(3-(benzo[d][1,3]dioxol-5-yl)-1H-pyrazol-5-yl)phenyl)amino)ethoxy)ethoxy)ethyl)amino)-3-(3,4-bis((4-fluorobenzyl)oxy)phenoxy)propan-2-ol) (50.0 g, 10.4%). (TLC: DCM/MeOH=10/1, $R_f$=0.6)

$^1$HNMR ($CDCl_3$, 400 MHz): δ 7.34-7.37 (m, 4H), 7.22-7.26 (m, 4H), 7.00-7.06 (m, 5H), 6.81 (d, *J* = 4 Hz, 1H), 6.77-6.79 (m 1H), 6.71 (s, 1H), 6.61-6.63 (m, 1H), 6.53 (d, *J* = 2.8 Hz, 1H), 6.33-6.36 (s, 1H), 5.96 (s, 2H), 4.97 (s, 2H), 4.95 (s, 2H), 4.33-4.35 (m, 1H), 3.89-3.98 (m, 2H), 3.64-3.73 (m, 8H), 3.37-3.40 (m, 2H), 2.93-2.99 (m, 4H)

LCMS [mobile phase: from 90% water (0.05% TFA) and 10% $CH_3CN$ to 5% water (0.1% TFA) and 95% $CH_3CN$ in 6.0 min, finally under these conditions for 0.5 min.] purity is >98% (254 nm), Rt = 3.069 min; Mass Calcd.:808.4; MS Found: 810.1 [MS+2].
